# Segregation of neural crest specific lineage trajectories from a heterogeneous neural plate border territory only emerges at neurulation

**DOI:** 10.1101/2021.08.08.455573

**Authors:** Ruth M Williams, Martyna Lukoseviciute, Tatjana Sauka-Spengler, Marianne E Bronner

## Abstract

The epiblast of vertebrate embryos is comprised of neural and non-neural ectoderm, with the border territory at their intersection harbouring neural crest and cranial placode progenitors. Here we profile avian epiblast cells as a function of time using single-cell RNA-seq to define transcriptional changes in the emerging ‘neural plate border’. The results reveal gradual establishment of heterogeneous neural plate border signatures, including novel genes that we validate by fluorescent *in situ* hybridisation. Developmental trajectory analysis shows that segregation of neural plate border lineages only commences at early neurulation, rather than at gastrulation as previously predicted. We find that cells expressing the prospective neural crest marker *Pax7* contribute to multiple lineages, and a subset of premigratory neural crest cells shares a transcriptional signature with their border precursors. Together, our results suggest that cells at the neural plate border remain heterogeneous until early neurulation, at which time progenitors become progressively allocated toward defined lineages.

## Introduction

During gastrulation, the ectoderm layer of the chordate embryo becomes segregated into the neural plate and the surrounding non-neural ectoderm. The neural plate ultimately generates the central nervous system (CNS), whereas the surrounding non-neural ectoderm forms the epidermis of the skin as well as the epithelial lining of the mouth and nasal cavities. At the interface of these tissues is a territory referred to as the ‘neural plate border’ which in vertebrates contains precursors of neural crest, neural and placodal lineages (Ezin et al., 2009, Streit, 2002). Neural crest and ectodermal placodes share numerous common features including the ability to migrate or invaginate and form multiple cell types. During neurulation, neural crest cells come to reside within the dorsal neural tube where they undergo an epithelial-to-mesenchymal transition (EMT). Subsequently they delaminate and migrate throughout the embryo, settle at their final destinations and differentiate into numerous derivatives including neurons and glia of the peripheral nervous system as well as cartilage, bone and connective tissue of the head and face. Like neural crest cells, cranial placode cells become internalised, and then differentiate into sensory neurons and sense organs (nose, ears, lens) of the head. Aberrant neural crest or placode development causes a number of developmental disorders affecting craniofacial structures (Siismets and Hatch, 2020, Vega-Lopez et al., 2018), the enteric nervous system (e.g. Hirschsprung’s disease) (Butler Tjaden and Trainor, 2013) and the heart (e.g., Persistent Truncus Arteriosus; CHARGE syndrome) (Pauli et al., 2017, Gandhi et al., 2020). Furthermore, a number of malignancies, including melanoma, neuroblastoma and glioma, are known to arise from neural crest derivatives (Tomolonis et al., 2018).

An ongoing question is whether individual neural plate border cells are specified toward a particular lineage (i.e., neural crest, placode or CNS) or if they have the potential to become any cell type that arises from the border. It has been suggested that progenitors of these different lineages may be regionalised within the neural plate border, with the more lateral cells contributing to the placodes and the more medial region giving rise to neural and neural crest cells (Schlosser, 2008). Such segregation has been proposed to result from the influence of graded expression of signalling factors emanating from surrounding tissues, e.g. Wnts and FGFs, on multipotent neural plate border cells (Schille and Schambony, 2017). Alternatively, individual neural plate border cells predetermined toward a particular lineage may be intermingled within the border. A recent study in the chick (Roellig et al., 2017) showed that while medial and lateral regions of the border can be discerned by lineage markers, there is significant co-expression of markers characteristic of multiple lineages (neural crest, neural plate and placodal) across this region. Moreover, this overlap of lineage markers within the neural plate border is maintained from gastrulation through neurulation suggesting that these cells may maintain plasticity through neurulation stages. Thus, while some neural plate border cells may be predisposed toward a particular fate, others retain the ability to generate multiple lineages. However, the timing at which neural plate border cells emerge and become distinguishable from neural and non-neural ectoderm has not been conclusively characterised, complicating the assessment of cell heterogeneity within the neural plate border territory.

One study using an *ex ovo* culturing method of explants from the chicken embryos proposed that a pre-neural plate border region is established as early as stage Hamburger and Hamilton (HH) 3 (Prasad et al., 2020). In addition, an *in vitro* model to derive neural crest cells from human embryonic stem cells shows that pre-border genes can be induced by Wnt signaling (Leung et al., 2016). However, these observations have not been thoroughly addressed *in vivo*. Therefore, the questions of how and when neural plate border cells establish/retain multipotency and the comprehensive gene dynamics underlying these processes remain open.

Single-cell RNA-sequencing (scRNA-seq) provides a unique platform to address the intriguing question of when a neural plate border transcriptional signature arises in an *in vivo* context. To this end, we examined single-cell transcriptomes from the epiblast of gastrulating to neurulating chick embryos to determine the time course of emergence of the neural plate border. The chick represents an ideal system for these studies since avian embryos develop as a flat blastodisc at the selected time points, highly reminiscent of human development at comparable stages. Interestingly, our results show that the border is not transcriptionally distinct until the beginning of neurulation (HH7), when neural plate border markers define a discrete subcluster of the ectoderm. Furthermore, RNA velocity measurements show that segregation of the neural plate border commences at HH6, but is more profoundly underway at HH7, with definitive neural crest clusters only emerging in the elevating neural folds. We further demonstrate that *Pax7*+ cells are not restricted to a neural crest fate and are capable of giving rise to all derivatives of the neural plate border. The data also reveal numerous novel factors dynamically expressed across the developing epiblast as well as revealing key drivers of neural plate border trajectories. Taken together, the data reveal dynamic changes in an emerging neural plate border that becomes progressively segregated into defined lineages, that is complete only late in neurulation.

## Results

### Single-cell analysis of the avian epiblast during gastrulation (HH4 – 5)

To resolve the transcriptional signatures of individual neural plate border cells in the context of the developing embryo, we first performed single-cell RNA-seq analysis of the epiblast of gastrulating chick embryos at stages HH4-5 (Hamburger and Hamilton, 1951). We used the Chromium 10X platform in order to recover a large number of cells, thus profiling the majority of neural plate border cells. As a reference for emergence of the neural plate border, we used *Pax7* and *Tfap2A* transcription factors, both of which are well-established early markers of the neural plate border (Basch et al., 2006, de Croze et al., 2011). *Tfap2A* demarcates the lateral aspect of the neural plate border from HH4 and is also expressed in the non-neural ectoderm (Figure 1A). *Pax7* is progressively enriched in the medial border region from HH5 (Figure 1D).

**Figure 1:**
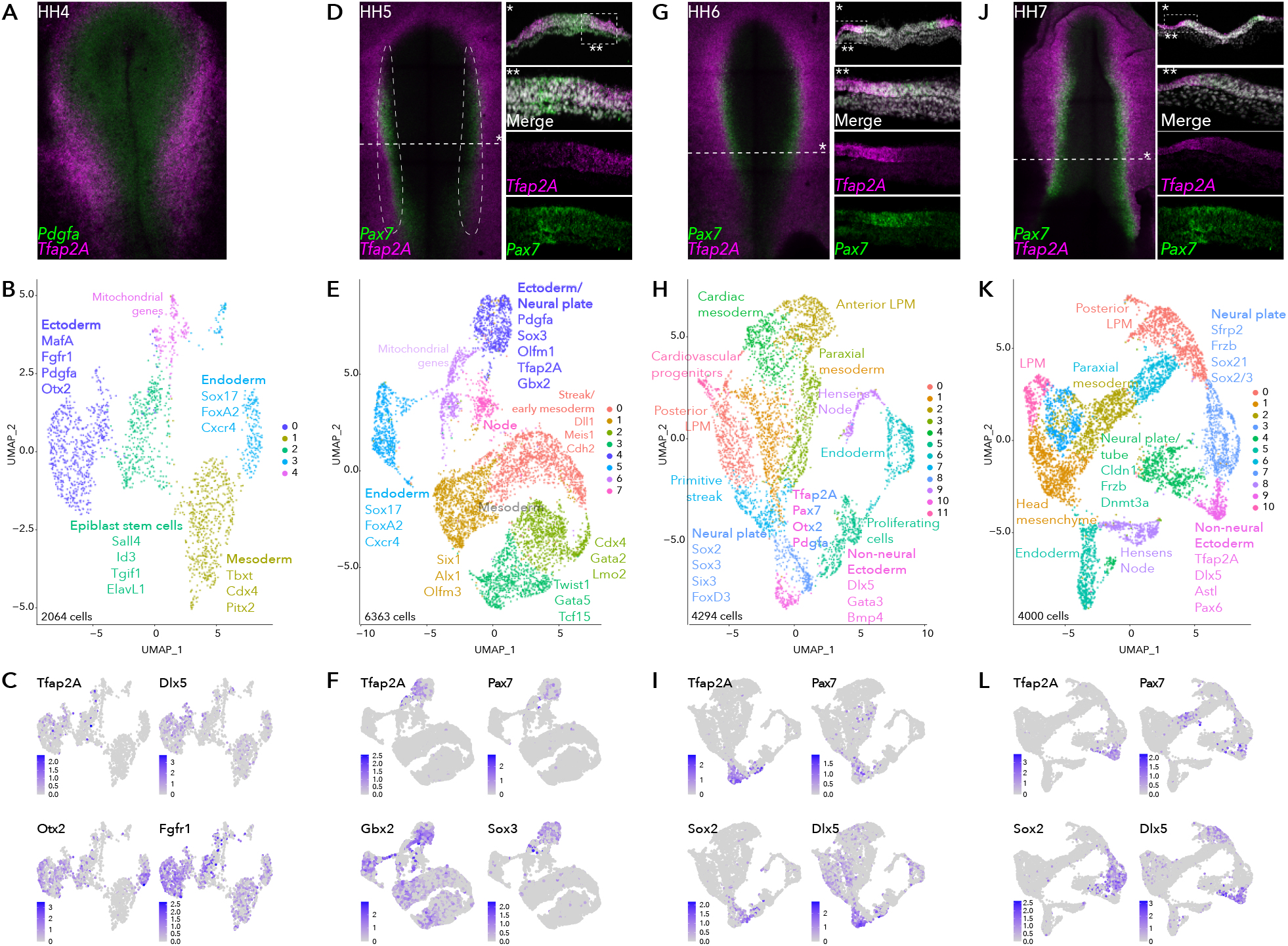
Single-cell RNA-seq of avian epiblast from HH4 through HH7. **(A)** *Tfap2A* expression at HH4, detected by HCR. **(B)** UMAP plot depicting 5 clusters resolved from 2064 epiblast cells at HH4. **(C)** Feature plots of selected genes in HH4 clusters. **(D)** *Pax7* and *Tfap2A* expression at HH5 detected by HCR. **(E)** UMAP plot depicting 8 clusters resolved from 6363 cells from dissected neural plate border regions at HH5, black dotted region in (E). **(F)** Feature plots of selected genes in HH5 clusters. **(G)** *Pax7* and *Tfap2A* expression at HH6 detected by HCR. **(H)** UMAP plot depicting 12 clusters resolved from 4294 epiblast cells at HH6. **(I)** Feature plots of selected genes in HH6 clusters. **(J)** *Pax7* and *Tfap2A* expression at HH7 detected by HCR. **(K)** UMAP plot depicting 11 clusters resolved from 4000 epiblast cells at HH7. **(L)** Feature plots of selected genes in HH7 clusters.

At HH4 (Figure 1A), we recovered 2398 cells from 8 embryos. Five distinct clusters (2064 cells) were resolved after quality control processing (Figure 1B, Figure S1A) and annotated by marker genes identified by single-cell differential expression (SCDE) analysis (Figure S1B/C). Focusing on the future neural plate border, two ectoderm clusters were recovered (HH4-Cl0, HH4-Cl2), identified by the enrichment in *Cldn1* expression (Figure S1C). HH4-Cl0 was enriched for neural plate markers (*Otx2* and *Fgfr1*), and featured low levels of *Tfap2A* and *Dlx5* (Figure 1C). Signaling molecules including *Sfrp2* and *Pdgfa* were also enriched in HH4-Cl0. *Sfrp2* and *Fgfr1* are expressed across the neural plate, extending to the neural plate border (Chapman et al., 2004, Lunn et al., 2007) and *Pdgfa* was expressed in the posterior epiblast (Yang et al., 2008). bHLH transcription factor *MafA*, more commonly associated with pancreatic beta-cell differentiation (Hang and Stein, 2011), but also reported in the developing chick neural plate (Lecoin et al., 2004), was enriched in HH4-Cl0 (Figure S1C). HH4-Cl2 was less distinctive but enriched in genes associated with pluripotency, such as *Sall4, Tgif1, ElavL1* (Lee et al., 2015, Ye and Blelloch, 2014, Zhang et al., 2006), (Figure S1B/C) suggesting these may represent residual epiblast stem cells.

To refine our analysis of the prospective neural plate border, we next performed 10X single-cell RNA-seq on dissected neural plate border regions from HH5 embryos (8 dissections) (Figure 1D) yielding 6363 cells and 6 clusters (Figure 1E, Figure S1D). Prospective neural plate border cells were confined to cluster HH5-Cl4, enriched for *Pax7* and *Tfap2A* expression, but also featuring a neural marker *Sox3*, and transcription factor *Gbx2*, which has a known role in neural crest induction in Xenopus (Li et al., 2009) (Figure 1F). While neural plate border markers were enriched in HH5-Cl4 cells, co-expression of neural and non-neural ectoderm factors here (Figure S1E/F) suggests this intermediate region is not yet distinguishable as a unique entity.

### Single cell analysis of the avian epiblast during neurulation (HH6 - 7)

During the process of neurulation in amniote embryos, the neural plate gradually folds inwards on itself, the border edges elevate forming the neural folds which, as neurulation progresses, fuse at the dorsal midline to form the neural tube. In the chicken embryo, this process is completed by HH8. Therefore, we performed single cell analysis at HH6 and HH7 to capture changes in the neural plate border as a function of time. While expression of *Pax7* was detectable but low during gastrulation stages (HH4/5), its expression is strongly enhanced by HH6/7 (Figure 1G/J), when this transcription factor marks more medial neural plate border cells.

Single-cell analysis at HH6 showed increased complexity revealing 12 distinct clusters from 4294 cells (Figure 1H, Figure S2A). A neural plate cluster (HH6-Cl8) was characterised by neural genes including *Sox2*/3, *Hes5* and *Otx2* (Figure 1I, Figure S2B/C) as well as neural crest regulators *FoxD3, Ednrb* and *Gbx2* (Figure S2B), whereas *Tfap2A* and *Dlx5* were enriched in the cluster HH6-Cl10. *Pax7* was heterogeneously detected at the interface of HH6-Cl8 and HH6-Cl10 (Figure 1I), indicating both these clusters harbour neural plate border cells.

At HH7 (Figure 1J), we identified 11 clusters from 4000 cells (Figure 1K, Figure S2D). Cells of the neural plate appeared in HH7-Cl3 and were distinct from the non-neural ectoderm cells found in HH7-Cl9 (Figure 1K/L). In addition to neural markers detected at earlier stages (*Otx2, Sox2, Sox3*), other neural/neural crest genes featured in HH7-Cl3, including *Zeb2* and *Zic2* (Figure S2E/F). Additional factors, such as *Pax6*, a crucial regulator of eye development (Liu et al., 2006) featured in the cluster HH7-Cl9 (Figure S2E). *Zfhx4* (Figure S2E), a zinc finger transcription factor previously observed at later stages in the neural crest and neural tube (Williams et al., 2019) was featured across HH7-Cl3 and HH7-Cl9 clusters.

Overall, across the stages analysed, we observed a progressive refinement of transcriptional signatures in individual ectoderm clusters reflecting neural versus non-neural lineages. While we identified markers of the neural plate border within multiple clusters, the border itself, surprisingly, is not distinguishable as a unique entity. Highlighting the heterogeneity of cells within the neural plate border, defined by combinatorial gene signatures shared with the surrounding neural plate and non-neural ectoderm.

### Subclustering reveals progressive transcriptional segregation of ectoderm cells

As the chick embryo undergoes gastrulation, ectodermal cells that will become neural, neural plate border, placode or epidermis remain in the upper layer of the embryo (epiblast), while mesoendodermal cells ingress and internalise at the primitive streak and Hensen’s node. To further resolve the transcriptional complexity of the developing neural plate border, we extracted and subclustered the ectodermal clusters at stages HH5, HH6 and HH7. These are designated HH5-Cl4, HH6-Cl8/10 and HH7-Cl3/9 (Figure 1E, H, K).

HH5-Cl4 resolved into 3 closely related clusters (Figure 2A). Neural plate (*Sox2, Otx2, Sox21, Six3*) subcluster (HH5-sub-1; Figure S3A) versus the non-neural ectoderm subcluster (*Tfap2A, Dlx5*; HH5-sub-2) were clearly delineated (Figure 2B). The third subcluster, HH5-sub-0, contained cells expressing caudal neural plate (*Msx1, Pdgfa, Wnt8A, Pou5f3*) and caudal epiblast genes (*Tbxt, Cdx2/4, Meis1/2*; Figure S3A/B). *Pax7*+ cells were most prominent in HH5-sub-0 (caudal) but a small portion of HH5-sub-2 (non-neural ectoderm) cells also expressed *Pax7* (Figure 2B). HH5-sub-0 cells co-expressed other neural plate border genes *Tfap2A, Dlx5, Msx1* and *Bmp4* with *Pax7*, these genes were also present in other clusters (Figure 2B).

**Figure 2:**
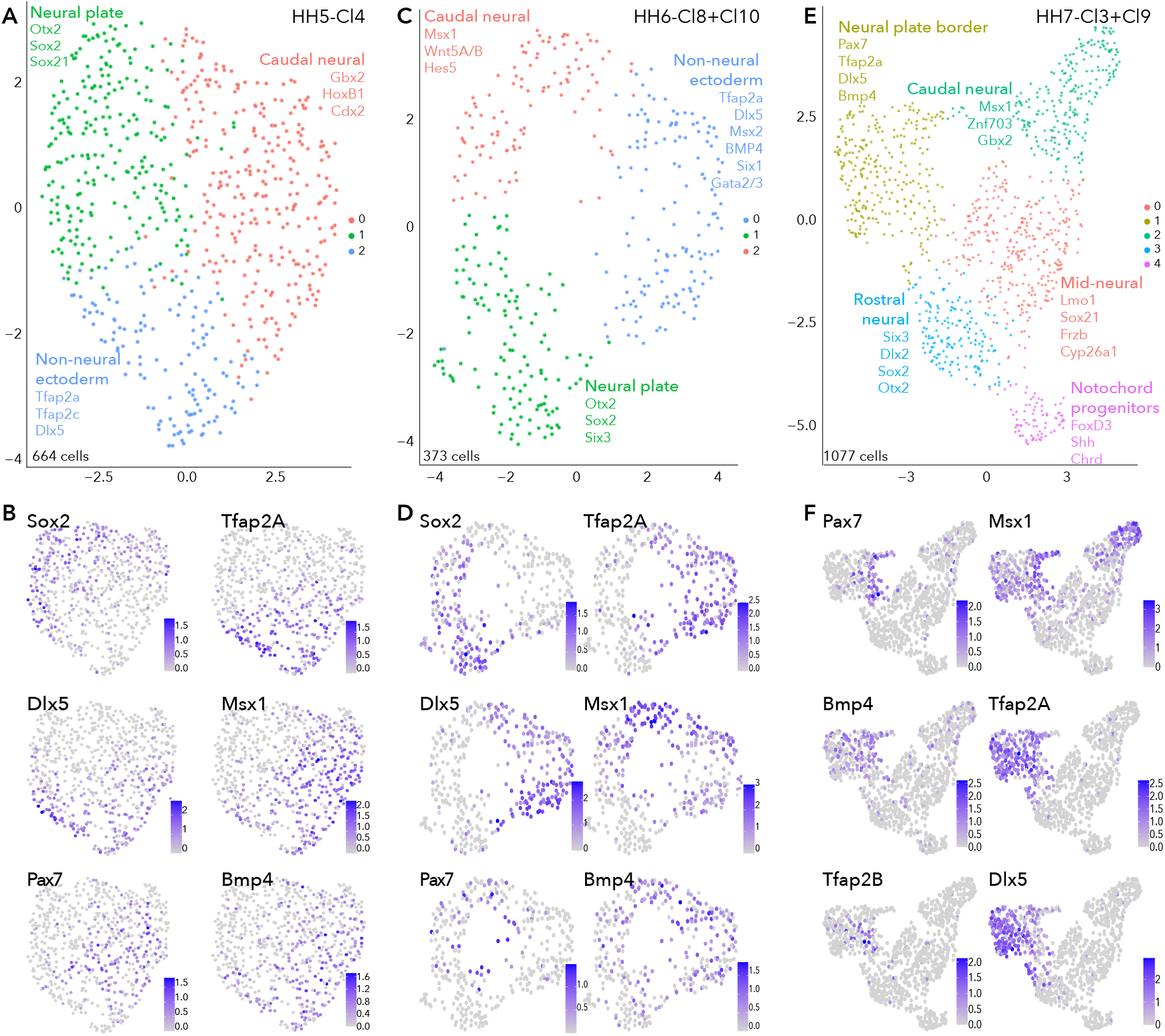
Subclustering ectoderm clusters extracted from whole epiblast data. **(A)** UMAP plot depicting 3 clusters resolved from HH5-Cl4. **(B)** Feature plots of selected genes in HH5 subclusters. **(C)** UMAP plot depicting 3 clusters resolved from HH6-Cl8 and HH6-Cl10. **(D)** Feature plots of selected genes in HH6 subclusters. **(E)** UMAP plot depicting 5 clusters resolved from HH7-Cl3 and HH7-Cl9. **(F)** Feature plots of selected genes in HH7 subclusters.

Similar signatures were found at HH6 (Figure 2C/D) but with increasing transcriptional segregation. Here, the neural plate was represented by HH6-sub-1 (*Sox2, Otx2, Six3*; Figure 2D, Figure S3C). Non-neural ectoderm markers (*Tfap2A, Dlx5*) were enriched in HH6-sub-0 (Figure 2D). *Pax7*+ cells clustered within the HH6-sub-2 (Figure 2D), together with caudal epiblast markers (*Tbxt, Cdx2/4*) (Figure S3C); however, *Pax7* expression also featured in HH6-sub-0 and HH6-sub-1, with *Pax7*+ cells positioned at the interface of the 3 subclusters (Figure 2D). These *Pax7*+ cells at the clusters’ interface also expressed *Tfap2A, Msx1, Dlx5* and *Bmp4* which were enriched in other clusters, as observed at HH5. This demonstrates that the neural plate border cluster is not yet distinct and indicates that neural plate border cells share signatures with other tissues at this stage (Figure 2D).

Further transcriptional refinements occurred between the HH5 and HH6 ectoderm subclusters. *Irf6* which was broadly present across HH5 subclusters, was now enriched in HH6-sub-0 (non-neural ectoderm), along with *Grhl3* which was present in a small portion of HH5-sub-2 (non-neural ectoderm; Figure S3B/D). *FoxD3* expression was barely detectable in cells of HH5-sub-1 (neural) but was clearly enriched in HH6-sub-1 (neural) (Figure S3B/D). *Gbx2* was broadly expressed across caudal epiblast and non-neural ectoderm subclusters (HH5-sub-0, HH5-sub-2 and HH6-sub-0, HH6-sub-2), while *Znf703* was restricted to more caudal epiblast cells (HH5-sub-0 and HH6-sub-2) (Figure S3B/D). *Gata2* and *Gata3* were found in a portion of HH5-sub-2 cells but their expression was expanded across HH6-sub-0 (Figure S3B/D). The placode marker *Six1* was found at low levels across all subclusters at HH5, but at HH6, *Six1* was restricted to only a portion of HH6-sub-0, potentially to cells within a placode progenitor niche (Figure S3B/D).

At HH7, ectodermal cells formed 5 discrete clusters (Figure 2E), suggesting lineage segregation was ongoing at this stage. Here neural plate border markers *Pax7, Msx1* and *Bmp4* were found in a niche of cells within HH7-sub-1, which broadly expressed *Tfap2A* and other non-neural ectoderm markers (Figure 2F, Figure S3E). Neural crest genes (*Tfap2B, Draxin, Snai2*) were arising in the *Pax7*+ domain (Figure S3F). HH7-sub-3 subcluster was characterised by neural markers; *Six3, Otx2, Sox21* and *Sox2* (Figure S3E). Some HH7-sub-1 cells also expressed placode markers (*Dlx5, Pax6, Six1/3*) but these cells did not co-express *Pax7* (Figure 2F; Figure S3E/F). Importantly, this reflects segregation of the neural plate border into medial (*Pax7*+) and lateral (*Tfap2A*+) regions, as well as highlighting the overlap and combination of genes co-expressed across these regions. HH7-sub-1 also featured *Irf6* expression, and *Grhl3* was present in a subset of cells which co-expressed *Irf6* and *Tfap2A* (Figure S3F). Another subset of HH7-sub-1 cells expressed *Pax6*, possibly already delineating prospective lens placode at this stage (Figure S3F), as these cells co-expressed *Six3*, known to activate *Pax6* by repressing Wnt signaling (Liu et al., 2006). HH7-sub-0 and HH7-sub-2 were also characterised as neural plate subclusters and could be resolved by axial markers, whereby mid-neural plate markers *Lmo1* and *Nkx6*.*2* were found in HH7-sub-0 and more caudally expressed genes *Cdx2/4, Hes5, Znf703* were found in HH7-sub-2 (Figure S3E/F). Cells in HH7-sub-4 were enriched for *Shh, Chrd* and *FoxD3*, suggesting they were notochord progenitors (Figure S3E).

In summary, analysis of our single cell HH5 and HH6 transcriptomes did not identify a distinct cell cluster marked by known neural plate border genes; rather cells expressing these markers were distributed across neural and non-neural clusters. Our analysis reveals the complex heterogeneity of the developing neural plate border and shows neural plate border cells are not distinct as a transcriptionally separate group of cells until HH7 when they also begin to segregate into presumptive medial and lateral territories. Moreover, many factors are shared between the neural plate border and other ectodermal cells as the neural plate border is progressively established from the neural plate and non-neural ectoderm.

### Validation of gene expression using Hybridisation Chain Reaction reveals dynamic in vivo expression patterns of novel neural plate border genes

We validated the *in vivo* expression pattern of intriguing genes identified in our single-cell datasets, using fluorescent /textitin situ hybridisation (Hybridisation Chain Reaction, HCR) (Choi et al., 2018), which enables simultaneous expression analysis of multiple transcripts. Embryos from HH4 through HH10 were screened to glean the time course of gene expression.

We identified several new genes in the ectoderm, many of which persisted into neural or non-neural tissues. At HH4 the chromatin remodeler, *Ing5*, Astacin-like metalloendopeptidase, *Astl*, and neuronal navigator 2, *Nav2* were enriched in the ectoderm cluster (HH4-Cl0) (Figure 3A, Figure S1C). At stages HH4-HH6 *Ing5* was expressed across the neural plate border, whereas *Astl* expression was predominantly detected in the posterior neural plate border (Figure 3B, Figure S4A). While both genes overlapped with *Tfap2A, Ing5* expression spread into the neural plate whereas *Astl* expression extended more laterally into the non-neural ectoderm (Figure 3B’). At HH8-*Astl* expression continued in the developing neural plate border and surrounding non-neural ectoderm, and also the neural plate (Figure 3C, 3C’). *Ing5* was also detected in the neural plate border and in the neural folds (Figure 3C). By HH8, *Astl* expression was prominent in the neural folds, most strongly in the posterior hindbrain (Figure 3D, 3D’). *Ing5* was detected along the neural tube at HH8 (Figure 3D, 3D’); however, by HH10 *Ing5* transcripts were no longer detectable. *Astl* expression was also broadly decreased at HH10, where activity was restricted to a small region of the neural tube in the hindbrain and the emerging otic placodes (Figure S4A). Consistent with these observations *Astl* and *Ing5* were detected in HH5-Cl4 and HH6-Cl10 (*Astl*)/HH6-Cl8 (*Ing5*). *Astl* was seen in HH7-Cl9, *Ing5* was not differentially expressed at this stage but was present in HH7-Cl3 (Figure S4B).

**Figure 3:**
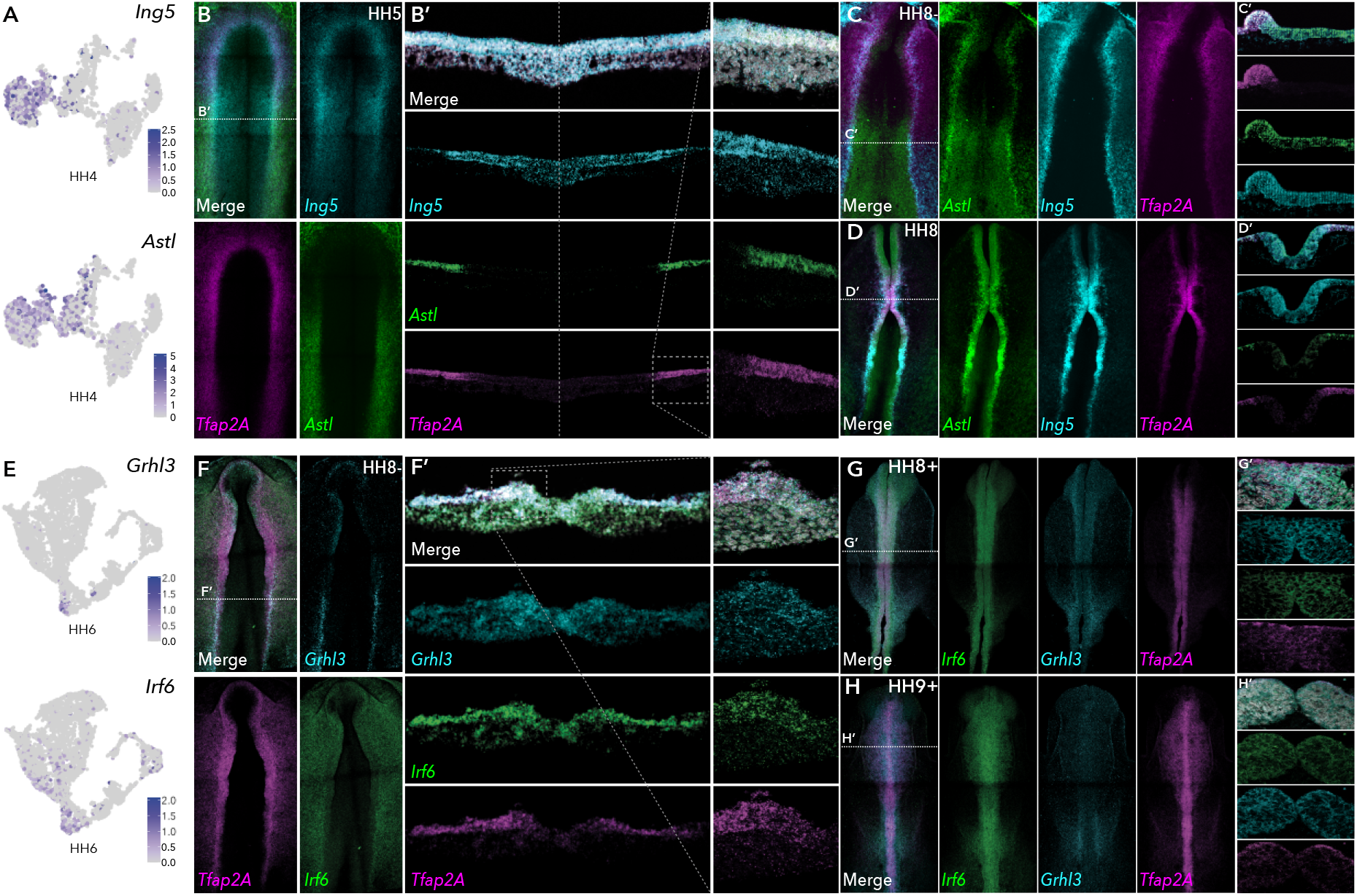
HCR validation of novel genes identified from whole epiblast data. **(A)** Feature plots of *Ing5* and *Astl* in HH4 data. **(B)** Whole mount HCR shows co-expression of *Ing5* and *Astl* with neural plate border marker *Tfap2A* at HH5. **(B’)** Transverse cryosection of (B) (dashed line). Dashed boxed region shown at high magnification in right panel. **(C-D)** Whole mount HCR shows co-expression of *Ing5* and *Astl* with neural plate border marker *Tfap2A* at HH8- **(C)** and HH8 **(D). (C’-D’)** Transverse cryosections of (C) and (D) respectively. **(E)** Feature plots of Grhl3 and Irf6 in HH6 data. **(F)** Whole mount HCR shows co-expression of Grhl3 and Irf6 with neural plate border marker *Tfap2A* at HH8-. **(F’)** Transverse cryosection of (F) (dashed line). Dashed boxed region shown at high magnification in right panel. **(G-H)** Whole mount HCR shows co-expression of Grhl3 and Irf6 with neural plate border marker *Tfap2A* at HH8+ (G) and HH9+ (H). **(G’-H’)** Transverse cryosections of (G) and (H) respectively.

*Nav2*, detected in ectoderm clusters HH4-Cl0, HH5-Cl4, HH6-Cl10 and HH7-Cl9 was expressed across the ectoderm and neural plate at HH5–HH7, where it over-lapped with the neural marker *Sox2*1 (Figure S4C/D), continuing into the neural folds at HH8 where *Nav2* was particularly prominent in the hindbrain (Figure S4C). At HH9 *Nav2* expression was restricted to rhombomeres 2 and 4, whereas *Sox2*1 was more broadly expressed along the neural tube axis (Figure S4C).

We identified a number of transcription factors in HH6-Cl8 (neural) and HH6-Cl10 (non-neural ectoderm). The latter was characterised by *Tfap2A* enrichment and also harboured *Grhl3* and *Irf6* (Figure 3E). *In situ* analysis showed *Grhl3* was specifically expressed in the neural plate border, most strongly in the posterior region at HH6 and HH8- (Figure S4E, Figure 3F). *Irf6* was expressed in the neural plate border as well as the surrounding non-neural ectoderm at HH6 and HH8- (Figure S4E, Figure 3F). Furthermore, we observed significant cellular co-expression of all three factors in the neural plate border (Figure 3F’). At HH8+, *Irf6* was restricted to the dorsal neural tube including premigratory neural crest cells, *Grhl3* was also detected here at lower levels but was present in the emerging otic placode (Figure 3G). By HH9+ *Grhl3* levels were broadly diminished but remained in the developing otic placode. *Irf6* was maintained in the neural tube during neural crest delamination and also enriched in the otic placode (Figure 3H).

In the ectodermal subclusters, we identified a number of novel factors including Wnt pathway genes. In HH6-sub-2, for example, we identified enrichment of Wnt signaling ligands *Wnt5A, Wnt5B, Wnt8A* (Figure S3C) as well as Wnt processing factors *Sp5* and *Sp8* (Figure 4A/E). Conversely Wnt antagonists *Sfrp*1, *Sfrp*2 and *Frzb* were enriched in HH6-sub-1 (Figure S3C). *Sp8* was identified in HH6-sub-2 (caudal neural), *in vivo* we found *Sp8* expression commenced from HH7 in the neural plate border, partially overlapping with *Tfap2A* (Figure 4B). The homeobox transcription factor, *Dlx6*, was present in HH7-sub-1, (neural plate, Figure 4A) and was expressed predominantly in the anterior neural plate border or pre-placodal region (Figure 4B). This pattern continued with higher levels of both *Sp8* and *Dlx6* at HH8- (Figure 4C). *Sp8* expression was restricted to the dorsal neural tube, whereas *Dlx6* spread more laterally (Figure 4C’). At HH9, *Sp8* was detected predominantly in the anterior neural tube but was also found in premigratory neural crest cells in the mid-brain region (Figure 4D, 4D’) but largely absent from the hindbrain region, though some expression was seen in the trunk premigratory neural crest (Figure 4D). *Dlx6* was also present in premigratory neural crest cells at HH9 (Figure 4D’), as well as in the lateral non-neural ectoderm (Figure 4D).

**Figure 4:**
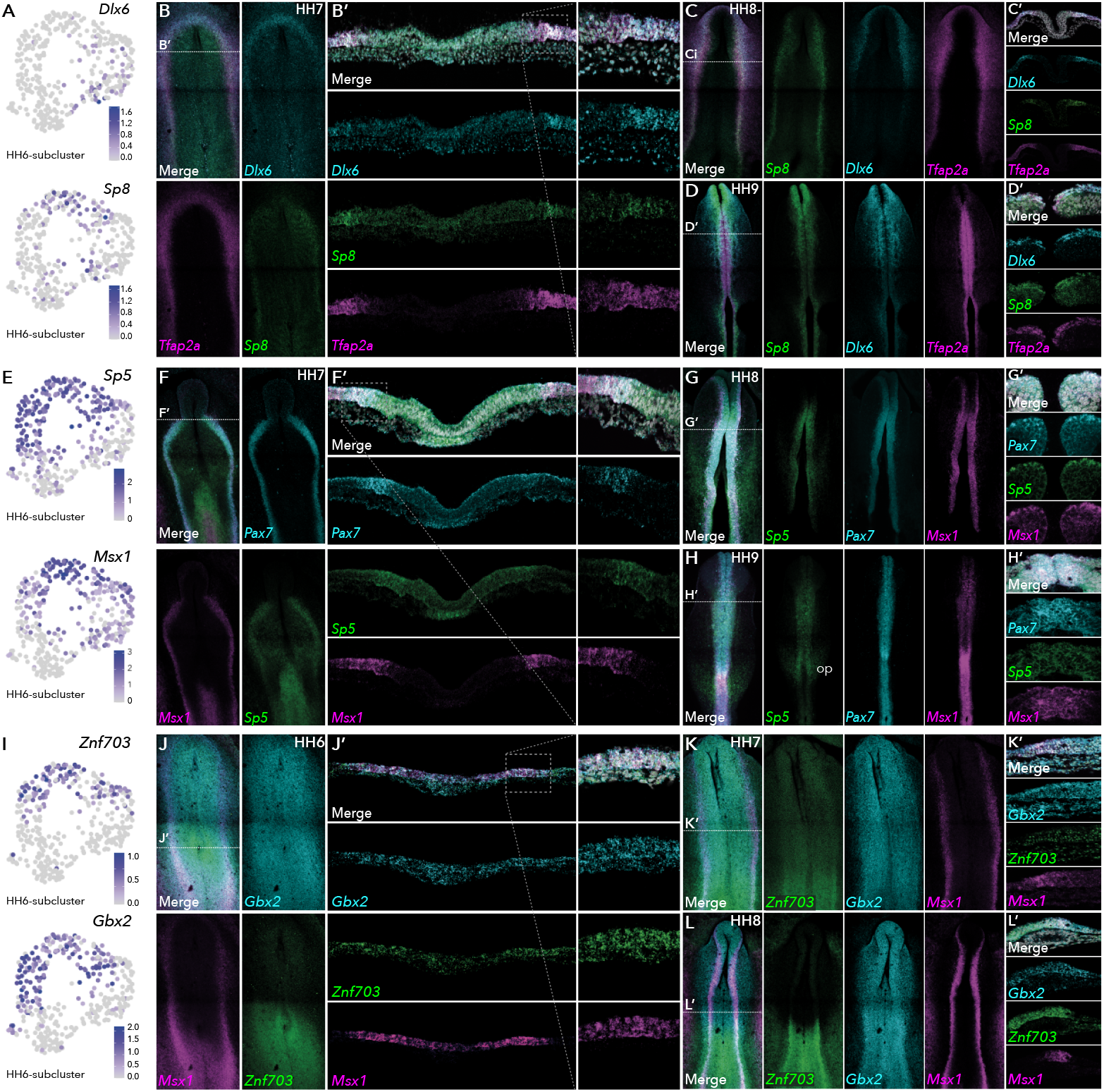
HCR validation of novel genes identified from extracted ectoderm clusters. **(A)** Feature plots of *Dlx6* and *Sp8* in HH6 subclusters. **(B)** Whole mount HCR shows co-expression of *Dlx6* and *Sp8* with neural plate border marker *Tfap2A* at HH7. **(B’)** Transverse cryosection of (B) (dashed line). Dashed boxed region shown at high magnification in right panel. **(C-D)** Whole mount HCR shows co-expression of *Dlx6* and *Sp8* with neural plate border marker *Tfap2A* at HH8- (C) and HH9 (D). **(C’-D’)** Transverse cryosections of (C) and (D) respectively.**(E)** Feature plots of *Sp5* and *Msx1* in HH6 subclusters. **(F)** Whole mount HCR shows co-expression of *Sp5* with neural plate border markers *Msx1* and *Pax7* at HH7. **(F’)** Transverse cryosection of (E) (dashed line). Dashed boxed region shown at high magnification in right panel. **(G-H)** Whole mount HCR shows co-expression of *Sp5* with neural plate border markers *Msx1* and *Pax7* at HH8 (G) and HH9 (H). **(G’-H’)** Transverse cryosections of (G) and (H) respectively. **(I)** Feature plots of *Znf703* and *Gbx2* in HH6 subclusters. **(J)** Whole mount HCR shows co-expression of *Znf703* and *Gbx2* with neural plate border marker *Msx1* at HH6. **(J’)** Transverse cryosection of (J) (dashed line). Dashed boxed region shown at high magnification in right panel. **(K-L)** Whole mount HCR shows co-expression of *Znf703* and *Gbx2* with neural plate border marker *Msx1* at HH7 (K) and HH8 (L). **(K’-L’)** Transverse cryosections of (K) and (L) respectively. Op; otic placode.

*Sp5* was enriched in HH6-sub-2 (Figure 4E) and HH7-sub-2 (Figure S3E). *Sp5* was detected in the anterior neural plate border where it overlapped with *Pax7* and *Msx1*, and in the posterior primitive streak/early mesoderm at HH7 (Figure 4F/F’). Later, (HH8-HH9) *Sp5* transcripts were restricted to the neural tube (Figure 4G/H) including premigratory neural crest cells (Figure 4G’/H’). At HH9 we also saw the onset of expression in the developing otic placodes (Figure 4H).

At HH6, HH6-Cl8 and HH6-Cl10 harboured *Znf703* and *Gbx2*, both of which were enriched in HH6-sub-2 (Figure 4I). *Znf703* has recently been demonstrated as RAR responsive factor important for neural crest development in Xenopus (Hong and Saint-Jeannet, 2017, Janesick et al., 2019). Consistent with this, we found *Znf703* expression in the posterior neural plate extending into the neural plate border from HH6 (Figure 4J/K). At HH8, *Znf703* transcripts also populated the posterior neural folds (Figure 4L) and partially colocalized with *Msx1* in premigratory neural crest cells (Figure 4L’). We found *Gbx2* broadly expressed across the epiblast from HH6 (Figure 4J/K), becoming more enriched in the neural folds by HH8 (Figure 4L). While both *Znf703* and *Gbx2* partially localized with *Msx1, Znf703* expression continued medially into the neural plate whereas *Gbx2* transcripts extended laterally into the non-neural ectoderm (Figure 4L’).

Neural clusters expressed numerous transcription factors including *Sox2*1 which was which was present in HH7-sub-0 and HH7-sub-3 (Figure S3E). *FezF2* together with *Otx2* and *Sox2* were also enriched in neural subclusters HH7-sub-3 (Figure S3C/E). *FezF2* has previously been shown to regulate Xenopus neurogenesis by inhibiting *Lhx2/9* mediated Wnt signaling (Zhang et al., 2014) Accordingly, *in vivo, FezF2* was expressed in the anterior neural folds at HH7 where it overlapped with *Otx2*, and in the neural tube at later stages (Figure S4F). *Lmo1* was also found in HH7-sub-0 (Figure S3E) and was expressed across the neural plate at HH6 through HH8, where it was also seen in the neural folds (Figure S4G).

### scVelo analysis shows developmental trajectories across the epiblast

The precise time at which the neural plate border segregates into the neural crest, placode and neural lineages has not been previously ascertained. To address this question, we used scVelo (Bergen et al., 2020) to resolve gene-specific transcriptional dynamics of these embryonic populations across time. This method is derived from RNA velocity (La Manno et al., 2018), which measures the ratio of spliced and unspliced transcripts and uses this information to predict future cell states, incorporating a critical notion of latent time that allows reconstruction of the temporal sequence of transcriptional steps. By generalising the RNA velocities through dynamic modelling, this method is ideally suited to our analysis, as it reflects transitions in progressing, non-stationary embryonic

Our whole embryo data demonstrated segregation of the germ layers at HH4 (Figure S5A) with all ectoderm cells (HH4-Cl0) aligning along the same trajectory. Epiblast stem cells (HH4-Cl2) were contributing to all other clusters (Figure S5A). At HH5, the majority of ectoderm cells (HH5-Cl4) still followed the same trajectory (Figure S5B); however, a portion of cells marked by *Tfap2A* and *Dlx5* were more transcriptionally stable (Figure S5C) indicating these are non-neural ectoderm cells whereas the differentiating population was marked by neural factors including *Zeb2* and *Cdh1*1 (Figure S5C).

A clear split in the ectoderm populations was first observed at HH6 with neural plate (HH6-Cl8) and non-neural ectoderm (HH6-Cl10) initiating separate trajectories (Figure 5A). *Sox2*+ cells contributed to the proliferating multipotent population. The remainder of HH6-Cl8 cells joined the posterior cell clusters with a discreet subset of cells engaged in a separate trajectory (Figure 5A). The *Sox2*+ subpopulation was specifically enriched for anterior neural markers *Otx2* and *Six3*, and pluripotency factor *FoxD3* (Figure 5B). In addition to the neural markers that characterised the rest of HH6-Cl8 (*Sox3, Hes5, Sfrp*2, *Frzb*) (Figure S2B/C), these cells also expressed neural plate border genes including; *Pax7, Tfap2A*, (Figure 1I), *Bmp4*, and *Msx1* (Figure 5B). Whilst these cells appeared to contribute to both neural and non-neural programmes, the low degree of splicing kinetics as compared to other subpopulations indicated they were relatively transcriptionally stable at this stage (Figure 5A).

**Figure 5:**
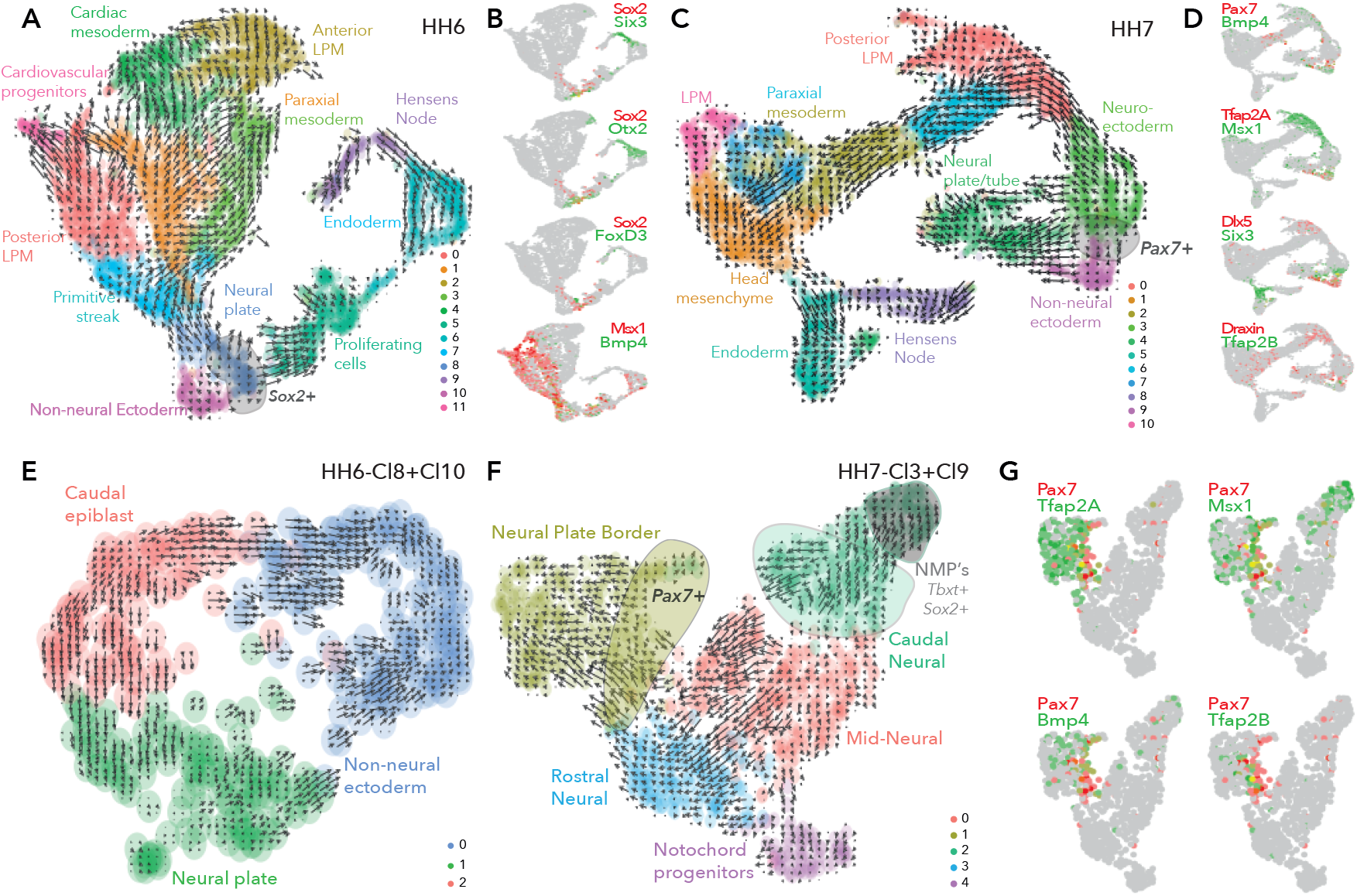
scVelo infers developmental trajectories across early epiblast stages. **(A)** UMAP embedding of clusters from whole epiblast data at HH6 showing predicted trajectories. **(B)** Colocalisation of selected genes across HH6 clusters **(C)** UMAP embedding of clusters from whole epiblast data at HH7 showing predicted trajectories. **(D)** Colocalisation of selected genes across HH7 clusters. **(E)** UMAP embedding of extracted ectoderm subclusters from HH6 showing predicted trajectories. **(F)** UMAP embedding of extracted ectoderm subclusters from HH7 showing predicted trajectories. **(G)** Colocalisation of selected genes across HH7 subclusters.

Interestingly, at HH7 we observed a convergence of trajectories between the neural plate (HH7-Cl3) and non-neural ectoderm (HH7-Cl9), likely representing the maturing neural plate border (Figure 5C). Within this population of cells, there was an overlap of neural plate border, ectodermal and placode gene expression (*Pax7, Bmp4, Tfap2A, Msx1, Dlx5* and *Six3*; Figure 5D) suggesting that they are not yet lineage-restricted but, rather, remain an intermingled heterogeneous progenitor population. Expression of early neural crest genes (*Draxin, Tfap2B*) also emerged in these converging cells (Figure 5D). Importantly, *Pax7* was exclusively found within this region (Figure 5D). Meanwhile, other cells from HH7-Cl3 and HH7-Cl9 were linking to the neural plate/tube trajectories (HH7-Cl4). We observed a portion of neural plate (HH7-Cl3) cells joining the posterior mesoderm cluster (HH7-Cl0), likely containing neuromesodermal precursors (NMPs) as suggested by the enrichment in *Tbxt* and *Sox2* expression (Figure S5D).

### Neural plate border lineages emerge progressively from the ectoderm over time

To further resolve the ectodermal trajectories, the extracted ectoderm clusters were modelled using scVelo. At HH5, each population (HH5-sub-0, caudal epiblast; HH5-sub-1, neural plate; HH5-sub-2, non-neural ectoderm) initiated a separate trajectory (Figure S5E). However, cells located centrally contributed to all 3 subclusters (Figure S5E) and expressed neural plate border genes including *Pax7* (Figure 2B).

At HH6 the trajectories were more distinct (Figure 5E). Caudal epiblast cells (HH6-sub-2) contributed to both neural plate and non-neural ectoderm populations. Within the neural plate cluster (HH6-sub-1) some cells were directed towards the non-neural ectoderm and others followed a separate trajectory. Likewise, the non-neural ectoderm cluster was subdivided between 2 parallel pathways with some mutual cross-over. Neural plate border cells found across all 3 clusters and at their interfaces (as depicted by *Pax7*; Figure 2D) largely joined the non-neural ectoderm clusters (Figure 5E).

HH7 trajectory map was more complex (Figure 5F). Here *Pax7*+ neural plate border cells (HH7-sub-1) projected toward all neural plate border derivatives, neural crest and placodes (HH7-sub-1) and neural tube (HH7-sub-0; Figure 5F). Some neural plate border cells appeared to be derived from the neural plate (HH7-sub-0/3), these cells expressed neural crest progenitor genes (*Tfap2B, Draxin, Snai2*,) (Figure 2F, Figure S3F). *Pax7*+ cells heterogeneously co-expressed other established neural plate border genes *Tfap2A, Msx1* and *Bmp4* (Figure 5G). This demonstrates that *Pax7*+ cells are capable of giving rise to multiple lineages and highlights heterogeneous combinations of factors at the neural plate border reflect multipotency. The remainder of HH7-sub-1 cells (*Pax7*-) were engaged in a separate trajectory (Figure 5F); these cells were enriched for *Dlx5* (Figure 2F), *Six1, Pax6* and *Gata3* (Figure S3F), suggesting a placodal fate. Overall, at HH7 scVelo analysis showed the neural plate border was beginning to segregate according to progenitor populations.

### Inferring developmental trajectories from presumptive ectoderm to premigratory neural crest cells

To assess the progression of ectodermal progenitors across time-points from epiblast to onset of neural crest emigration, we first integrated the ectoderm clusters from each stage namely; HH5-Cl4, HH6-Cl3+9 and HH7-Cl8+10. Here we also included 10X scRNA-seq data obtained from premigratory neural crest cells (HH8+) from our previous work (Williams et al., 2019), such that we could trace trajectories from the neural plate border to the *bona fide* neural crest.

Combining these datasets revealed that the non-neural ectoderm from both HH6 and HH7 grouped together, as did the neural plate clusters from the same stages (Figure 6A). This suggests successful inter-sample incorporation into a single reference with no resultant batch bias (Figure S5F). Ectoderm cells from HH5 were spread amongst the neural plate and the non-neural ectoderm clusters from HH6 and HH7. The neural crest clusters were more discrete, with the *bona fide* neural crest cluster (*FoxD3, Tfap2B, Sox10*) clearly segregated from neural (*Sox3, Pax6, Cdh2*) and mesenchymal (*Pitx1, Twist1, Sema5B*) neural crest clusters as previously described (Williams et al., 2019). The HH8+ dataset also included a neuroepithelial-like cluster, characterised by *Cdh1, Epcam* and *FoxG1* expression. Interestingly these cells grouped with the non-neural ectoderm clusters from HH6 and HH7, including neural plate border cells (*Pax7, Tfap2A*; Figure 6A/B), suggesting early neural crest cells retain properties of their predecessors.

**Figure 6:**
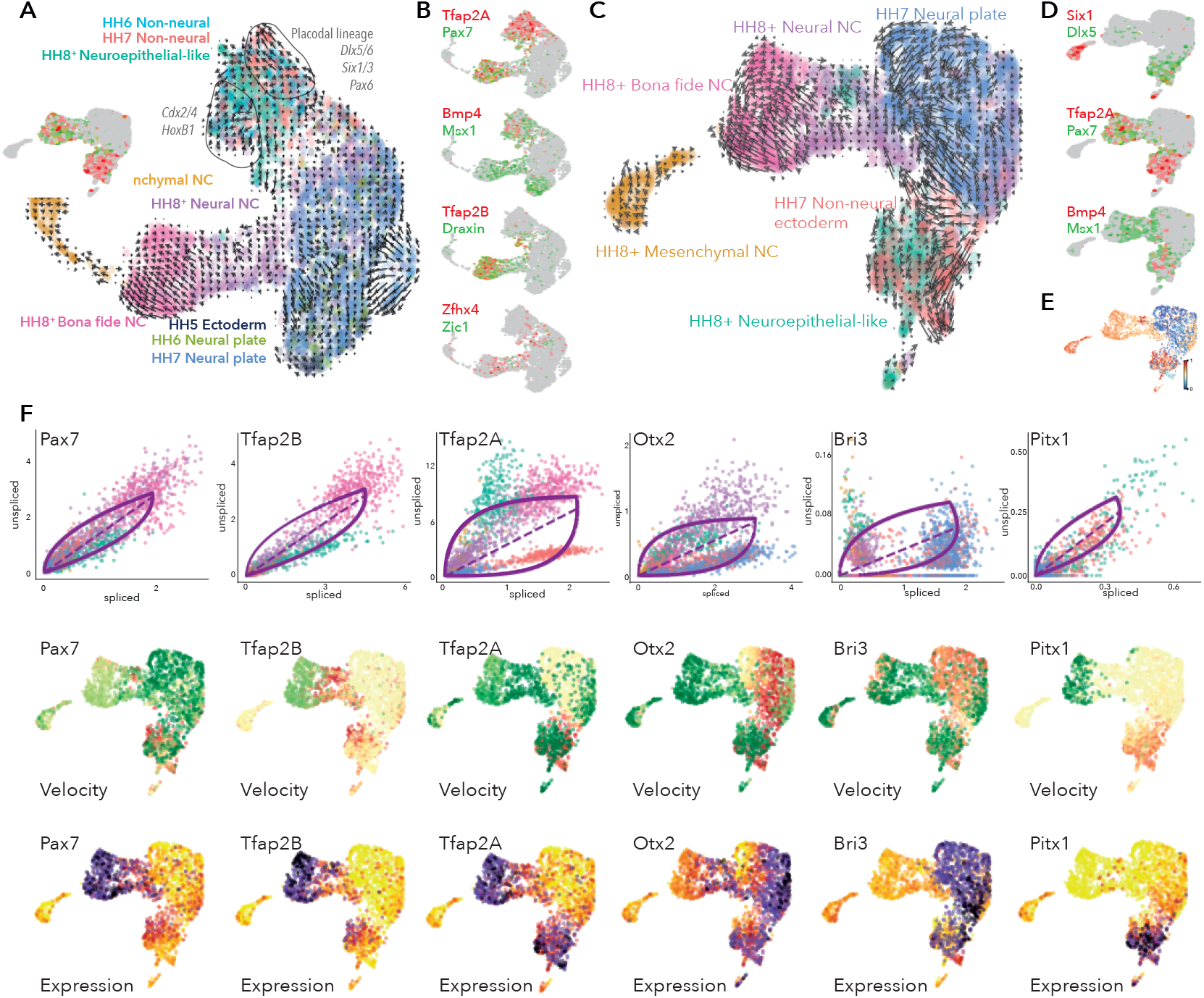
Developmental trajectory analysis from the neural plate border to premigratory neural crest cell. **(A** UMAP embedding of ectoderm clusters combined with premigratory neural crest data showing predicted trajectories. **(B)** Colocalisation of selected genes across combined data set. **(C)** UMAP embedding of HH7 ectoderm clusters combined with premigratory neural crest data showing predicted trajectories. **(D)** Colocalisation of selected genes across HH7/premigratory neural crest combined data set. **(E)** Latent time data from HH7/premigratory neural crest combined dataset. **(F)** Phase portraits (top row), velocity plots (middle row) and expression plots (bottom row) of selected genes across the HH7/ premigratory neural crest combined data sets. NC; neural crest.

scVelo showed that non-neural ectoderm cells from HH6/HH7 were split between two trajectories (Figure 6A). Placodal markers (*Dlx5*/6, *Six3* and *Pax6*) defined one distinct trajectory (Figure S5E), whereas the other trajectory was less well defined but could be distinguished by posterior epiblast markers (HoxB1, *Cdx4*) (Figure S5H). The neuroepithelial-like cells from HH8+ were emerging between these trajectories where they were joined by other cells from HH6/HH7 non-neural ectoderm clusters. These cells were enriched for *Pax7* and other neural plate border markers *Tfap2A, Msx1, Bmp4* and (Figure 6B). Early neural crest genes including *Tfap2B* and *Draxin* were also emerging here (Figure 6B). While these cells had less defined trajectories, they were generally directed towards the neural-neural crest lineage (Figure 6A). Neural plate border and neural crest markers also extended into a heterogeneous region of cells, where a significant portion of the neural-neural crest and neural plate cells were highly integrated, suggesting some cells from these populations shared transcriptional signatures. This region was further defined by *Zfhx4* and *Zic1* (Figure 6B) which are expressed in the neural plate border, neural plate and premigratory neural crest cells (Khudyakov and Bronner-Fraser, 2009, Williams et al., 2019). Some neural-neural crest cells within this region appeared to be directly derived from the neural plate.

Since we did not see evidence of neural crest lineages until HH7, we assessed the developmental trajectories from HH7 ectoderm clusters directly to *bona fide* neural crest cells (HH8). To this end, we extracted the ectoderm clusters (HH7-Cl3+Cl9) and combined these cells with the premigratory neural crest data set from HH8+ embryos and used scVelo to plot the trajectories across these cells/stages. Analysis of the pooled dataset yielded clearly defined developmental trajectories splitting from the HH7 ectoderm clusters, whereby the non-neural ectoderm cells (HH7-Cl9) gave rise to the neuroepithelial-like cells from the HH8+ dataset, and the neural plate (HH7-Cl3) cells were contributing directly to the neural-neural crest (Figure 6C). Non-neural ectoderm cells were also initiating a separate trajectory, likely representing placodal lineages as indicated by *Six1, Dlx5* (Figure 6D). While *Tfap2A* was expressed across the non-neural ectoderm cluster, *Pax7* was found in just a subset of non-neural ectoderm cells and in neuroepithelial-like cells, where *Tfap2A* was also found. Both factors were also enriched in *bona fide* neural crest, though *Pax7* was also detected in the neural-neural crest (Figure 6D). Other neural plate border markers (*Bmp4, Msx1*) were found in the non-neural ectoderm and some neuroepithelial-like cells, as well as extending into the interface of neural plate and neural-neural crest populations (Figure 6D).

Taken together, combining our 10X data-sets from all stages reinforced the notion that the neural plate border territory is not specified to a neural crest fate until HH7 since scVelo analysis on the entire dataset (including HH5 and HH6 data) generated ambiguous and non-contiguous developmental trajectories (Figure 6A). While neural crest trajectories were not clearly defined until HH7, placodal trajectories could be discerned from HH6. Latent time analysis corroborated these observations, indicating the progression of transcriptional maturity and corresponding differentiation status across the HH7 non-neural ectoderm cells into the neuroepithelial-like cells and, similarly, from the neural plate cells into the neural-neural crest and canonical neural crest populations (Figure 6E). Furthermore, the data showed a sub-population of premigratory neural crest cells shared transcriptional signatures with their progenitor cells.

### Analysis of dynamic transcriptional trajectories during lineage restriction reveals key lineage specific drivers

We next sought to determine key genes driving the observed developmental trajectories in the context of neural crest specification from the neural plate border. Dynamical modelling of transcriptional states within the HH7/8 combined data-set revealed a number of highly ranked dynamic genes including *Pax7* and *Tfap2B. Pax7* splicing increased progressively from the non-neural ectoderm cells to the neuroepithelial-like cells and lower splicing was observed in neural crest cells where the transcripts stabilized such that increased expression was observed (Figure 6F). Splicing of *Pax7* transcripts was also increased in the putative placodal lineage from the non-neural ectoderm (Figure 6F). *Tfap2B* splicing was not evident in the HH7 data, but was increased in the neuroepithelial-like cells and stabilized in neural crest populations (Figure 6F). *Tfap2A* showed more complex velocities, whereby high levels of spliced transcripts were detected in non-neural ectoderm cells from HH7 likely driving these cells towards the placode trajectory (Figure 6F). We also observed a progressive increase in *Tfap2A* splicing across the neuroepithelial-like cells to the *bona fide* neural crest. This suggested that by HH8, *Tfap2A* transcripts are stable in the neuroepithelial-like population but upregulated in the *bona fide* neural crest, consistent with the observed expression dynamics (Figure 6F). Another neural plate border gene, *Bmp4*, was dynamically regulated in non-neural ectoderm but downregulated in the neural plate (Figure S5I).

We also identified a number of more novel factors putatively involved in ectoderm lineage progression. *Otx2* was driving neural plate cells towards the neural-neural crest cluster, consistent with previous findings from functional perturbation studies (Williams et al., 2019). However, *Otx2* also seemed to be driving non-neural ectoderm cells towards the neuroepithelial-like population (Figure 6F). *Lmo1* was also driving the neural plate cells towards the neural-neural crest; however, unlike *Otx2, Lmo1* expression was not maintained in the neural crest populations (Figure S5I).

*Bri3* was identified as a highly ranked dynamic gene and was upregulated in neural plate cells and some non-neural ectoderm cells. Down-regulation of *Bri3* in neural crest cells suggested an early role in ectoderm lineage trajectories for this novel factor (Figure 6F). *Nav2* and *Sox11* were both highly expressed in the neural plate and downregulated in the non-neural ectoderm. *Sox11* was also expressed in the mesenchymal neural crest cluster where it was highly spliced, potentially representing a dual segregated role for this factor in both neural plate and mesenchymal crest lineages (Figure S5I). *Nav2* velocities increased progressively from the non-neural ectoderm to the neural plate; consistently, a subset of non-neural ectoderm cells joined a trajectory with the neural plate cells (Figure S5I) where *Nav2* splicing was highest (Figure S5I). We found *Pitx1* and *Dlx6* to be dynamically active across the merged HH7/8 data-set, potentially driving non-neural ectoderm to the neuroepithelial-like population (Figure 6F, Figure S5I).

This analysis provides insight into potential candidates driving lineage specific circuits underlying the progressive segregation of the early ectoderm into neural and non-neural ectoderm and concomitantly initiating the emergence of early neural crest and placode progenitors.

## Discussion

Here we use single-cell RNA-sequencing of chick embryos from late gastrula through early neurula to characterise the development of the neural plate border and its derivatives, the neural crest and cranial placode precursors. Our data show that the neural plate border, as defined by co-expression of *Tfap2A* and *Pax7* first emerges at HH5, but is not fully transcriptionally defined until HH7. Previous work has pointed to the presence of a pre-border region at blastula stages harbouring neural crest progenitors demarcated by *Pax7* expression (Basch et al., 2006; Prasad et al., 2020). However, this was observed in explanted cultures. In contrast, we do not detect significant *Pax7* expression until HH5. This suggests that cells within the explants may not have been fully specified at the time of explantation but cell interactions coupled with autonomous programmeming enabled the cells to continue their specification programme.

We identified several genes in our dataset that were not previously known to function in the neural plate border. For example, *Irf6* has a well-established role in craniofacial development (Fakhouri et al., 2017, Ingraham et al., 2006, Wang et al., 2003) but has yet to be explored at earlier stages of development. *Grhl3* has been shown to work in a module with *Tfap2A* and *Irf6* during neurulation (Kousa et al., 2019). In addition, Wnt signaling pathway genes like *Sp5* and *Sp8* were prominent. Wnt signaling has a well-established role in neural plate border specification; for example, Wnt signals are required to activate the earliest neural crest genes *Tfap2A, Gbx2* and *Msx1. Sp8* is known to play a role in processing Wnt signals during limb development (Haro et al., 2014, Kawakami et al., 2004), but has also been described in neural patterning and craniofacial development (Sahara et al., 2007, Zembrzycki et al., 2007), whereby loss of *Sp8* in mice caused severe craniofacial defects due to increased apoptosis and decreased proliferation of neural crest cells (Kasberg et al., 2013). However, an early role for *Sp8* has not been explored. *Sp5* has recently been implicated in the neural crest gene regulatory network, where it was found to negatively regulate *Axud1* to help maintain neural crest cells in a naïve state (Azambuja and Simoes-Costa, 2021). Therefore, *Sp5* and *Sp8* may represent hitherto unknown components of Wnt mediated induction of neural plate border lineages. *Gbx2* was recently reported as the earliest Wnt induced neural crest induction factor in Xenopus. Perturbation of *Gbx2* inhibited neural crest development while the placodal population was expanded (Li et al., 2009). Interestingly, our analysis reveals distinct heterogeneity of Wnt signaling factors, indicating differential cellular responses to Wnt signaling within the developing neural plate border, which may contribute to driving different lineage trajectories.

We used scVelo analysis of transcriptional and splicing dynamics across clustered and aligned single-cell transcriptomes to observe the dynamics of developmental trajectories during ectoderm lineage segregation. This enabled us to follow the transcriptional velocity of individual genes and resolve their dynamics at the single-cell level. Significantly this provides a high-resolution temporal dimension to our understanding of the changing ontology of the neural plate border and its derivatives. The results suggest that the ectoderm is initiating the division of trajectories between neural and non-neural progenitors at HH5, but the neural plate border is not yet transcriptionally distinct. At HH6 and HH7 the neural plate border population is emerging at the interface of these tissues, from which multiple lineages manifest by HH7. This analysis also shows neural-neural crest cells are the first to surface from the neural plate border and neural plate. From these, the *bona fide* neural crest emerges followed by mesenchymal neural crest. Furthermore, this analysis shows a subset of premigratory neural crest cells share signatures with their precursors from neural plate border cells from HH6 and HH7, demonstrating that some early neural crest cells are not yet restricted to a particular lineage.

Taken together our data show that cells of the emerging neural plate border are not characterised by unique transcriptional signatures, but share features with cells of the surrounding ectoderm. This highlights the heterogeneity of the neural plate border whereby *Pax7*+ cells are integrated with other ectoderm populations. While *Pax7* is the predominant factor in the neural plate border and has been suggested to label neural crest precursors as early as HH5 (Basch et al., 2006), the complexity and co-expression signatures are what endow neural plate border cells with their unique multipotency compared to neural plate and ectoderm cells. Interestingly, we note that neural plate border signatures are not apparent until HH7, later than previously suggested. By contrast, placodal trajectories were discernible from HH6. Moreover, we find *Pax7*+ cells give rise to all neural plate border lineages but lose multipotency signatures at the onset of lineage specific trajectories. Our analysis provides important insights into genes underlying the progressive segregation of the emergence of early neural crest and placode progenitors at the neural plate border. By revealing lineage trajectories over developmental time, this resolves the timing of neural plate border lineage segregation whilst also informing on dynamics of multipotency programmemes.

## Supplementary results

HH4: Other clusters at HH4 were readily identified as mesoderm (HH4-Cl1) or endoderm (HH4-Cl3) as characterised by the expression of *Cdx4* and *Pitx2* for mesoderm and *Sox17, FoxA2* and *Cxcr4* for endoderm (Figure 1B, Figure S1B/C. HH4-Cl2 was enriched for genes associated with pluripotency including *Sall4, Tgif1, ElavL1* (Lee et al., 2015, Ye and Blelloch, 2014, Zhang et al., 2006) (Figure S1B/C). Mitochondrial genes and cell migration factors (*Cxcl12, Itgb1, Tgfbr1*) were enriched in HH4-Cl4 suggesting these are highly active migrating cells (Figure 1B, Figure S1C). scVelo analysis revealed that mesoderm (HH4-Cl1) and endoderm cells (HH4-Cl3) were following distinct trajectories as individual populations (Figure S5A). At HH5, mesoderm cells (HH5-Clusters 0-3) were segregating and endoderm cells (HH5-Cl5) were also maintaining their distinct trajectory (Figure S5B).

HH6: Mesodermal cells were found in several clusters: anterior lateral plate mesoderm (HH6-Cl2; *Pitx2, Alx1*); posterior lateral neural plate (HH6-Cl0; *Gata2, HoxB5*). Cardiac mesoderm markers were enriched in HH6-Cl4 (*Tcf21* and *Gata5*) and HH6-Cl11 (*Lmo2, Ets1, Kdr*). HH6-Cl1 and HH6-Cl3 represented paraxial mesoderm (*Msgn1, Mesp1*). Endoderm cells formed HH6-Cl6 (*Sox17, FoxA2*). Hensen’s node and primitive streak markers (*Dll1, Fgf8, Noto, Chrd*) were identified in HH6-Cl9 and HH6-Cl7 respectively. We also detected a cluster of cells (HH6-Cl5) with high levels of mitochondrial genes (*Cox1/3*), this cluster was also enriched for factors associated with highly proliferative cells (*Pdia3, Igf1r*) (Figure S2C).

HH7: Aspects of the mesoderm were discernible across several clusters (HH7-Cl0, 1, 2, 6, 7, 10). HH7-Cl5 was defined by endoderm markers and HH7-Cl8 represented Hensen’s node and the primitive streak (Figure S2F).

## Author contributions

Conceptualisation, R.M.W., T.S.S., M.E.B; Methodology, R.M.W. Software, R.M.W., M.L., Validation, R.M.W.; Formal Analysis, R.M.W., M.L.; Investigation, R.M.W.; Writing R.M.W.; Writing – Review & Editing, all authors; Visualisation R.M.W., T.S.S., M.E.B; Supervision, T.S.S., M.E.B; Funding Acquisition, M.E.B.

## Acknowledgements

Fluorescence activated cell sorting was performed at California Institute of Technology Flow Cytometry Facility using BD Biosciences FACSAria IIu Cell Sorter with Patrick Cannon. 10X libraries were prepared in the Thomson lab at California Institute of Technology with assistance from Jeff Park. Illumina sequencing was performed at the Millard and Muriel Jacob at California Institute of Technology with Igor Antoshechkin. HH4 10X library was constructed at MRC Weatherall Institute of Molecular Medicine, University of Oxford with assistance from Kevin Clark (FACS facility), Dr Neil Ashley (single-cell facility) and Tim Rostron (NGS sequencing facility). Confocal microscopy was performed within the Biological Imaging Facility at the Beckman Institute, California Institute of Technology with assistance from Dr Giada Spigolon. We thank members of the Bronner and Sauka-Spengler labs for their support and helpful discussions. This work was funded by NIH R01DE027538 to MEB.

## Materials and Methods

### Chick embryos

Fertilized chicken eggs, obtained from Sunstate Ranch Sylmar CA, were incubated at 37°C with approximately 40% humidity. Embryos were staged according to (Hamburger and Hamilton, 1951).

### Preparing embryos for FAC-sorting

Appropriately staged embryos were extracted using the filter paper based ‘easy-culture’ method. Eggs were opened after desired incubation period, albumin was removed and embryos were lifted from the yolk using punctured filter paper, this procedure is described in detail elsewhere (Williams and Sauka-Spengler, 2021b). Embryos were kept in Ringers solution and dissected to remove all extra-embryonic material. Embryos were then dissociated for fluorescence activated cell sorting (FACS) as previously described (Williams and Sauka-Spengler, 2021a). Embryos were processed by FACS using 7-AAD as a live/dead stain such that healthy individual cells were obtained with a reliable cell count.

### 10X single-cell RNA-seq library preparation and sequencing

Approximately 10,000 cells/stage were collected by FACS into 2ul Hanks buffer then loaded onto the 10X Genomics Chromium platform. Single-cell RNA-seq libraries were generated using the Chromium Single Cell 3’ Library and Gel Bead Kit v3 (10X Genomics, Cat. #1000075) (HH4 data was obtained using v2 chemistry) as per the manufacturers protocol. Libraries were quantified using Qubit (Life Tech Qubit high sensitivity DNA kit Cat. #Q32854) and Kapa (Kapa Biosystems, KAPA Library Quantification Kit, Cat. #KK4835). scRNA-seq libraries were sequenced on Illumina HiSeq2500 platform in rapid run mode with on-board clustering and sequencing depth of 300 million reads. The run type was: paired end 28(read1)-8(index)-91(read2). HH4 10X scRNA-seq library was sequenced on Illumina NextSeq500 platform using high output v2.5 150-cycle kit in 26 x 8 x 0 x 98 mode.

### 10X single-cell RNA-seq data analysis

Fastq files were generated using cellranger v3.1.0 mkfastq (Zheng et al., 2017). Cellranger was also used for base calling, demultiplexing and mapping to the galgal6 genome assembly. A custom galgal6 genome was constructed using the mkref function whereby selected 3’UTRs were extended according to manually annotation. From HH4 embryos 2398 cells were recovered with 144,605 mean reads/cell, 1836 median genes/cell, 6180 median UMI counts/cell and 15,370 total genes. From HH5 embryos 6723 cells were recovered with 44,343 mean reads/cell, 3659 median genes/cell, 14,229 median UMI counts/cell and 18,916 total genes. From HH6 embryos 4553 cells were recovered with 30,466 mean reads/cell, 3396 median genes/cell, 13,309 median UMI counts/cell and 17,828 total genes. From HH7 embryos 6585 cells were recovered with 20,044 mean reads/cell, 2696 median genes/cell, 8821 median UMI counts/cell and 17,873 total genes. Count matrices were generated using the cellranger count function and exported to R-studio for downstream analysis using Seurat-v3 (Stuart et al., 2019). Matrices were filtered to remove barcodes with fewer than 500 genes and more than 3500-5500 genes (Figure S1/S2) and high mitochondrial content (*>*0.5%). UMI counts were normalised and following principal component analysis linear dimension reduction was conducted (HH4; resolution 0.2, dims 1:20, HH5; resolution 0.4, dims 1:15, HH6; 0.6, dims 1:20, HH7; resolution 0.4, dims 1:20). Clustered cells were visualised on a UMAP plot and differentially expressed genes were identified. Ectoderm clusters were extracted and re-clustered using the ‘subset’ command, analysis was performed as for whole embryo data.

### scVelo analysis

The analysis was performed using the scVelo dynamical modelling pipeline (Bergen et al., 2020) in python. scVelo analysis on combined datasets from different stages: HH5, HH6, HH7 and HH8 datasets were integrated using Seurat v3 by finding inter-sample anchors followed by integration. UMI counts and mitochondrial transcript percentage were regressed out during integrated object scaling, 30 npcs were used for dimension reduction to plot a single reference UMAP. For downstream scVelo analysis HH5, HH6, HH7 and HH8 ectodermal/pro-neural clusters or HH7 and HH8 ectodermal/pro-neural clusters were subtracted from the integrated dataset, followed by rescaling and PCA reduction using 10 or 9 dimensions respectively to plot UMAPs. Loom files containing spliced /unspliced transcript expression matrices were generated for each individual stage using velocyto.py pipeline (La Manno et al., 2018). Loom cell names were renamed to match Seurat object cell names and only Seurat-filtered cells were selected for trajectory analysis. Loom files from individual stage samples were merged using loompy.combine function. Seurat generated UMAP coordinates, clusters and cluster or stage colours were added to the filtered loom cells.

### Hybridisation chain reaction

Fluorescent in situ hybridsation chain reaction was performed using the v3 protocol (Choi et al., 2018). Briefly, embryos were fixed in 4% paraformaldehyde (PFA) for 1 hour at room temperature, dehydrated in a methanol series and stored at −20°C at least overnight. Following rehydration embryos were treated with Proteinase-K (20mg/mL) for 1-2.5 minutes depending on stage (1 min HH4-6, 2.5 mins for older embryos) at room temperature and post-fixed with 4% PFA for 20 min at room temperature. Embryos were washed in PBST for 2x 5 min on ice, then 50% PBST / 50% 5X SSCT (5X sodium chloride sodium citrate, 0.1% Tween-20) for 5 min on ice and 5X SSCT alone on ice for 5 min. Embryos were then pre-hybridized in hybridisation buffer for 5 min on ice, then for 30 min at 37°C in fresh hybridization buffer. Probes were prepared at 4pmol/mL (in hybridization buffer), pre-hybridization buffer was replaced with probe mixture and embryos were incubated overnight at 37°C with gentle nutation. Excess probes were removed with probe wash buffer for 4x 15 min at 37°C. Embryos were pre-amplified in amplification buffer for 5 min at room temperature. Hairpins were prepared by snap-cooling 30pmol (10ml of 3mM stock hairpin) individually at 95°C for 90 s and cooled to room temperature for minimum 30 min, protected from light. Cooled hairpins were added to 500µl amplification buffer. Pre-amplification buffer was removed from embryos and hairpin solution was added overnight at room temperature, protected from light. Excess hairpins were removed by washing in 5X SSCT 2x 5 min, 2x 30 min and 1x 5 min at room temperature. Embryos were mounted on slides and imaged using Zeiss LSM 880 Upright confocal microscope. Images were processed using Zeiss Zen software, Z-stacks scans were collected at 6µm intervals across approximately 70-100µm, maximum intensity projections of embryo z-stacks are presented. Tile scanning was used (2×3) and stitched using bidirectional stitching mode, with overlap of 10%. For sectioned samples, images were obtained on a Zeiss LSM 880 Upright confocal with 20X and 40X oil immersion objectives, single z-slices are shown.

### Cryosectioning

Following HCR and whole mount imaging, selected embryos were prepared for cyrosectioning Embryos were placed in a 15% sucrose solution at 4°C overnight then 15% sucrose / 7.5% Gelatin overnight at 37°C. Embryos were transferred to 20% gelatin and incubated at 37°C for 4 hr then mounted in 20% gelatin, snap frozen in liquid nitrogen and stored at −80°C. Sections were taken at 10µm intervals.

## Supplemental Material

**Supplementary figure S1.**
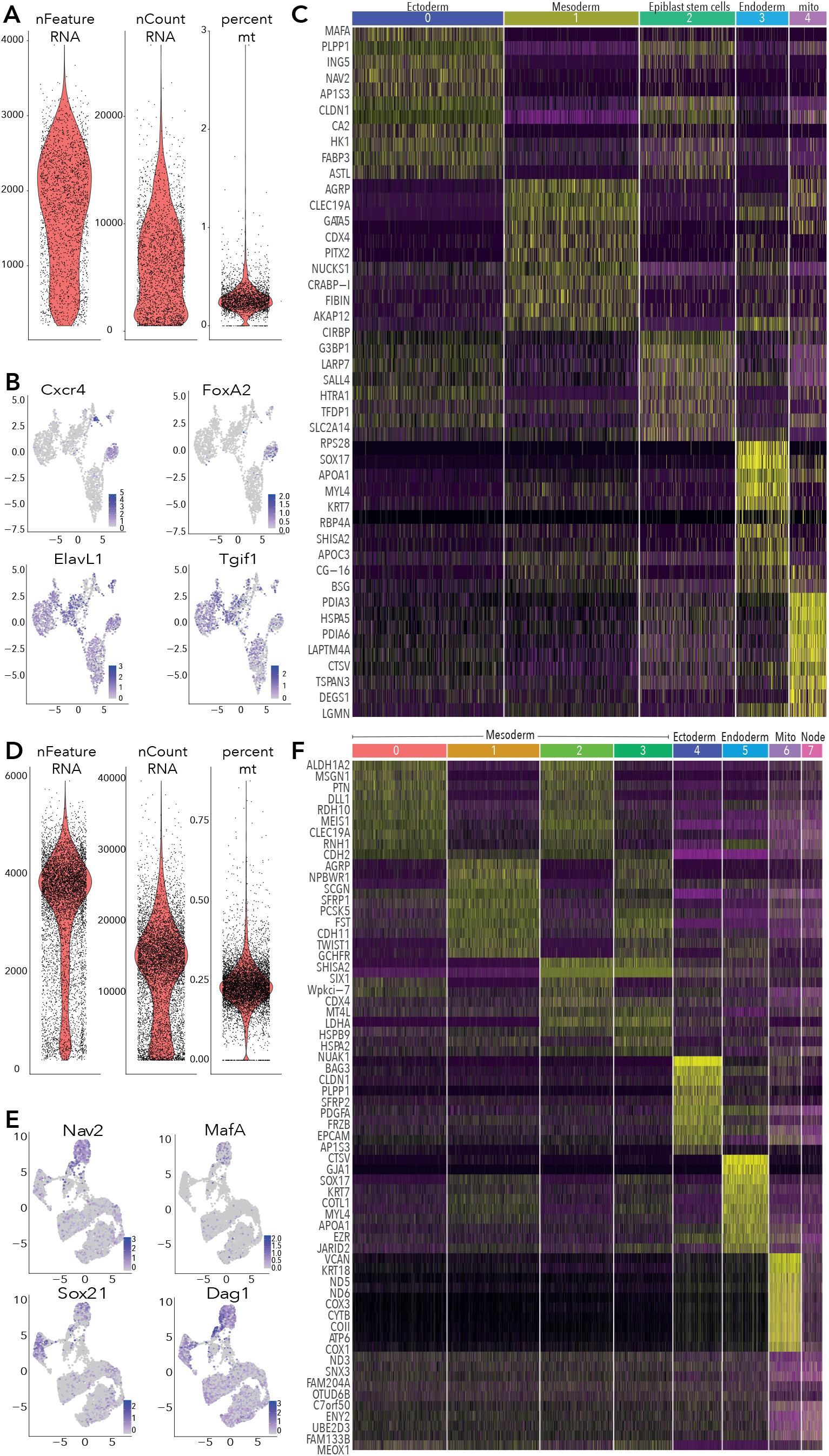
Quality control and supporting data for HH4 and HH5 10X data. (A) Violin plots showing RNA features, counts and percentage of mitochondrial reads per cell from HH4 data. Cells were filtered to exclude those with *<*500 and *>*3500 features (genes) and *>*0.5% mitochondrial gene content. (B) Feature plots of selected genes in HH4 clusters. (C) Heatmap depicting top 10 differentially expressed genes across each cluster from HH4. (D) Violin plots showing RNA features, counts and percentage of mitochondrial reads per cell from HH5 data. Cells were filtered to exclude those with *<*500 and *>*5500 features (genes) and *>*0.5% mitochondrial gene content. (B) Feature plots of selected genes in HH5 subclusters. (C) Heatmap depicting top 10 differentially expressed genes across each cluster from HH5.

**Supplementary figure S2.**
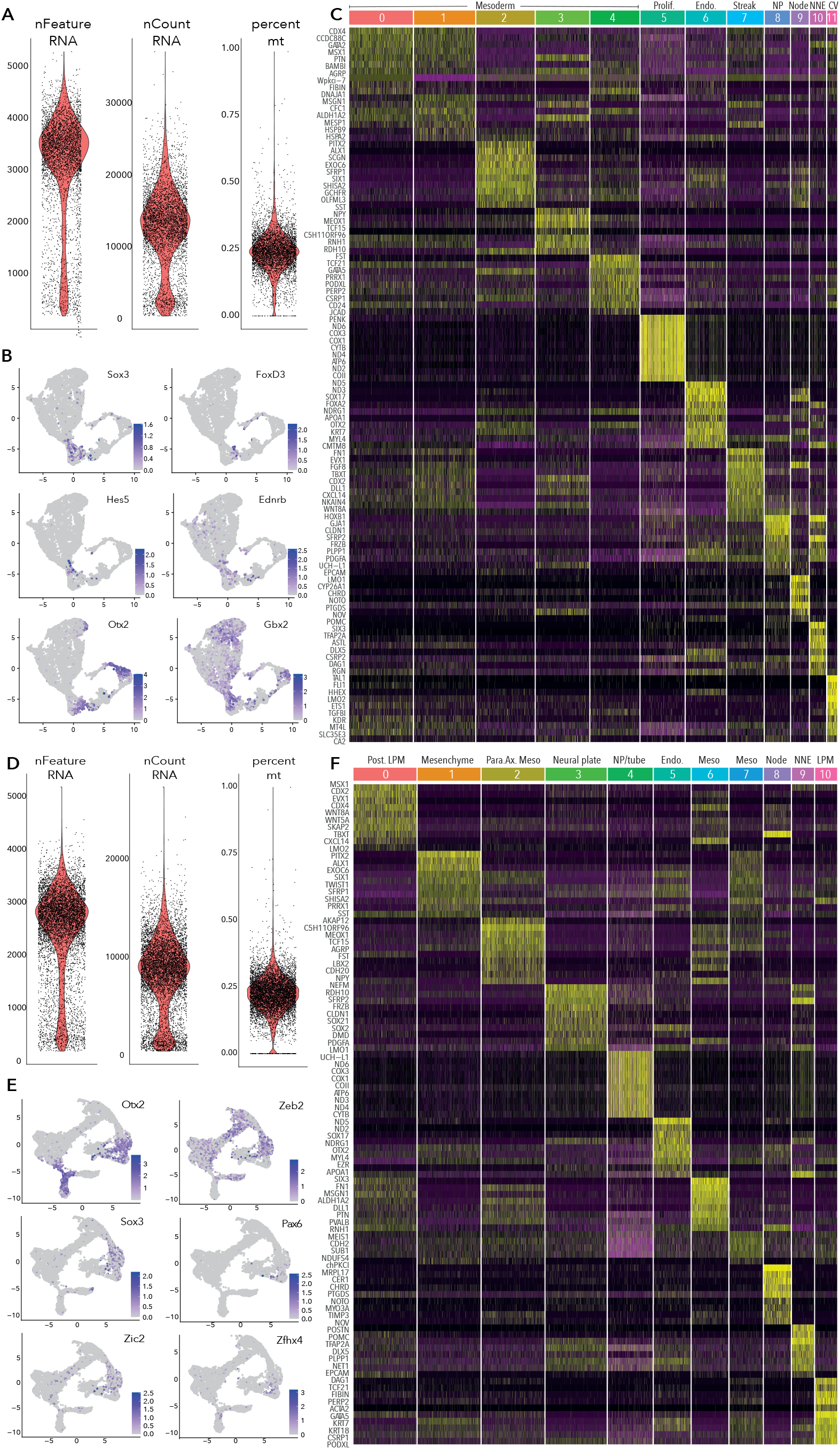
Quality control and supporting data for HH6 and HH7 10X data. (A) Violin plots showing RNA features, counts and percentage of mitochondrial reads per cell from HH6 data. Cells were filtered to exclude those with *<*500 and *>*5000 features (genes) and *>*0.5% mitochondrial gene content. (B) Feature plots of selected genes in HH6 clusters. (C) Heatmap depicting top 10 differentially expressed genes across each cluster from HH6. (D) Violin plots showing RNA features, counts and percentage of mitochondrial reads per cell from HH7 data. Cells were filtered to exclude those with *<*500 and *>*4000 features (genes) and *>*0.5% mitochondrial gene content. (B) Feature plots of selected genes in HH7 clusters. (C) Heatmap depicting top 10 differentially expressed genes across each cluster from HH7.

**Supplementary figure S3.**
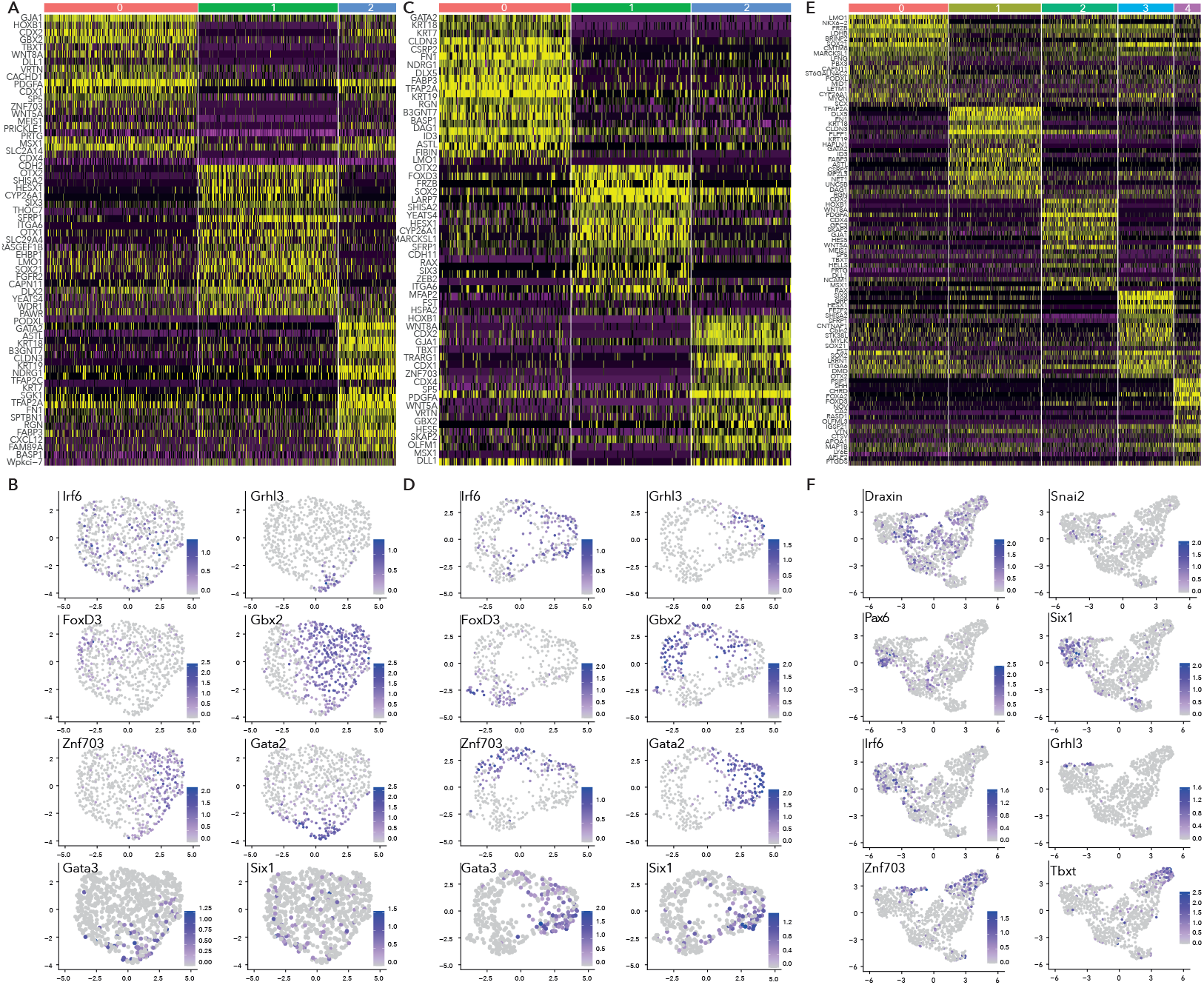
Supporting data for subclustering HH5, HH6 and HH7 ectoderm clusters. (A) Heatmap depicting top 20 differentially expressed genes across HH5 subclusters. (B) Feature plots of selected genes in HH5 subclusters. (C) Heatmap depicting top 20 differentially expressed genes across HH6 subclusters, (D) Feature plots of selected genes in HH6 subclusters. (E) Heatmap depicting top 20 differentially expressed genes across HH7 subclusters. (F) Feature plots of selected genes in HH7 subclusters.

**Supplementary figure S4.**
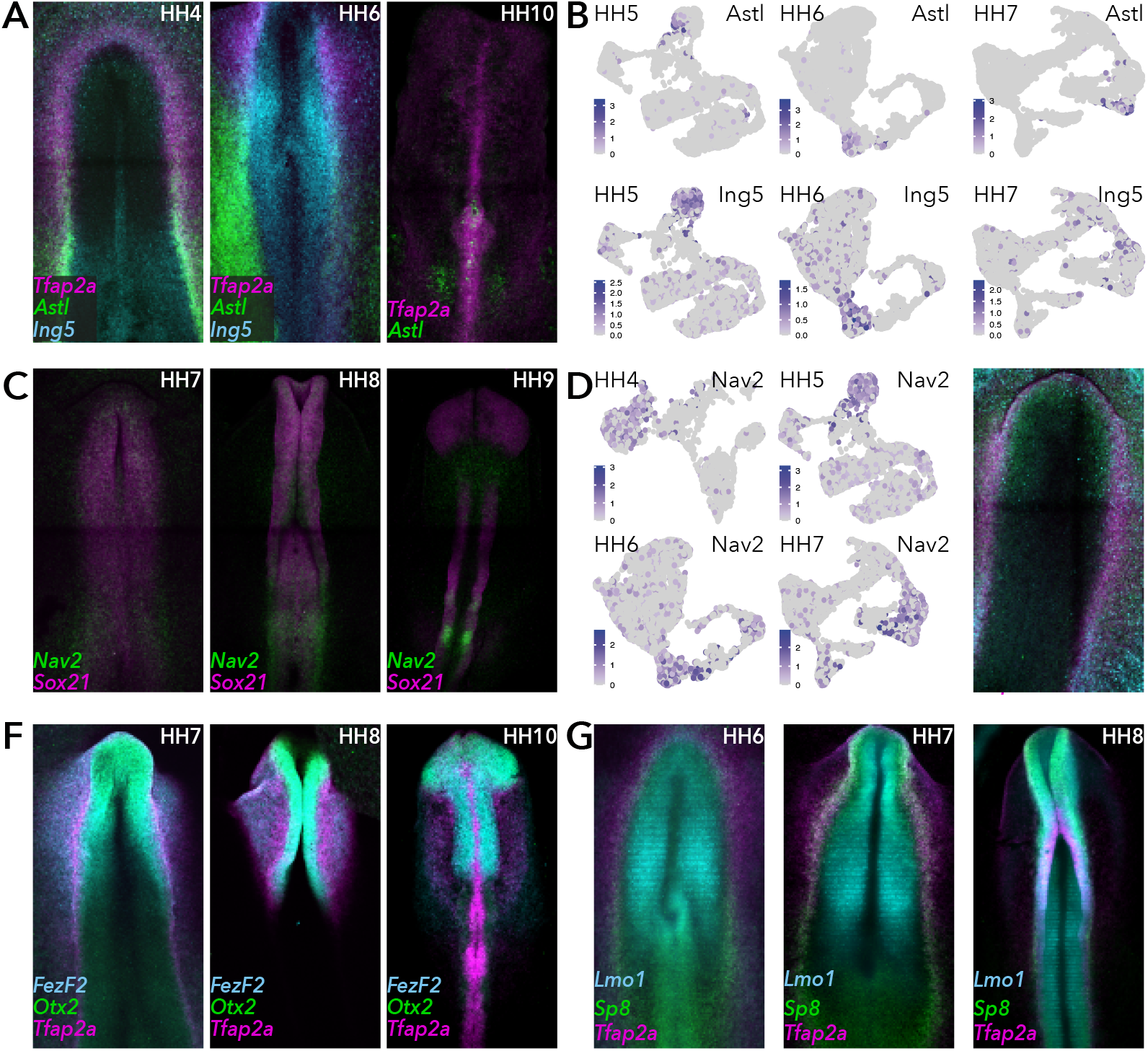
HCR validation of selected genes. (A) Whole mount HCR shows co-expression of *Astl* and *Ing5* with neural plate border marker *Tfap2A* at stages HH4, HH6, HH10. (B) Feature plots of *Astl* and *Ing5* across whole epiblast data sets at stages HH5, HH6 and HH7. (C) Whole mount HCR shows co-expression of *Nav2* and *Sox21* at stages HH7, HH8 and HH9. (D) Feature plots of *Nav2* across whole epiblast data sets. (E) Whole mount HCR shows co-expression of *Grhl3, Irf6* and *Tfap2A* at HH7. (F) Whole mount HCR shows co-expression of *FezF2* and *Otx2* with neural plate border markers *Tfap2A* at stages HH7, HH8 and HH10. (G) Whole mount HCR shows co-expression of *Lmo1* and *Sp8* with neural plate border markers *Tfap2A* at stages HH6, HH7 and HH8.

**Supplementary figure S5.**
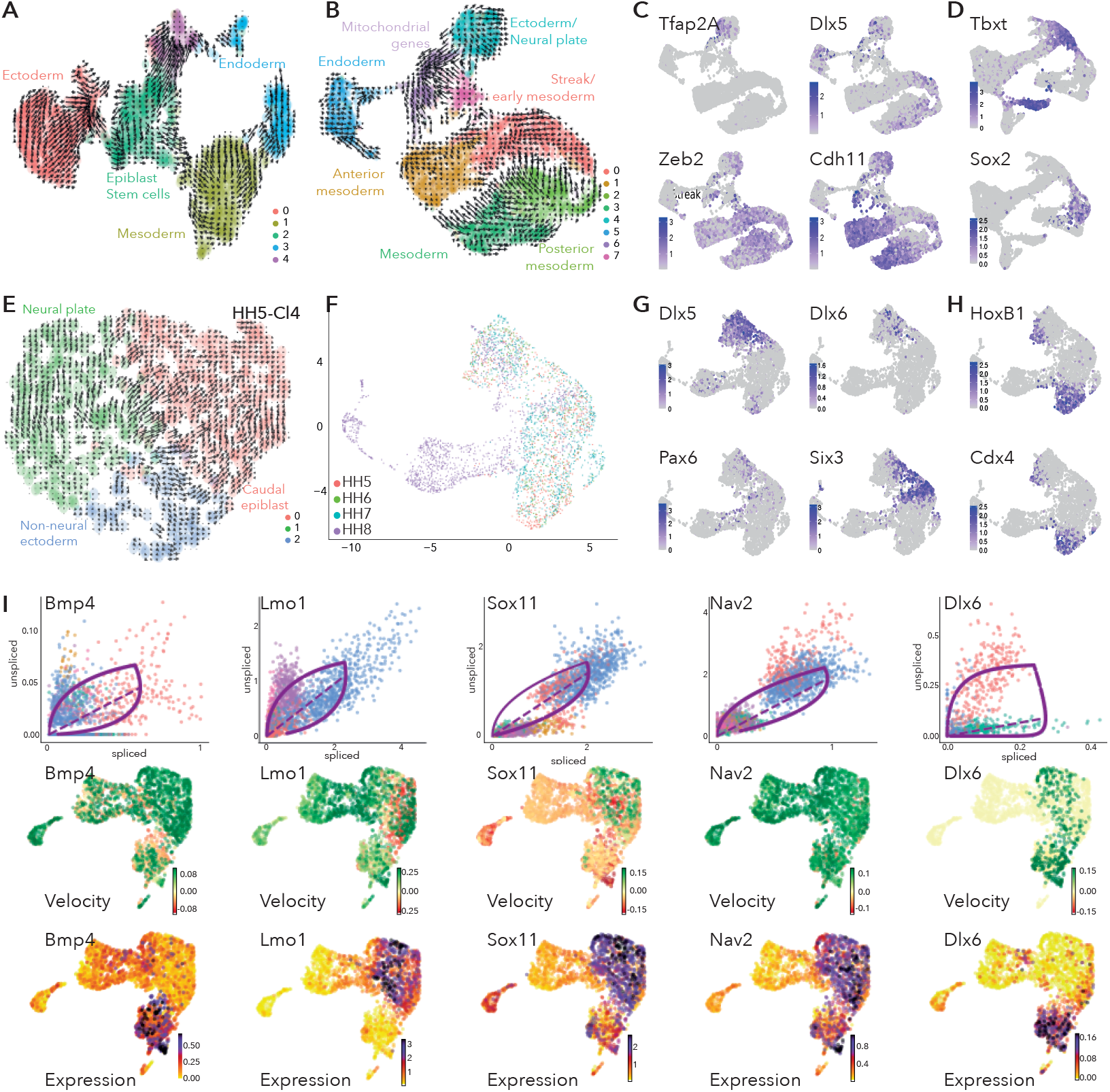
scVelo analysis at HH4 and HH5 and supporting data for scVelo analysis at HH6 and HH7. (A) UMAP embedding of clusters from whole epiblast data at HH4 showing predicted trajectories. (B) UMAP embedding of clusters from whole epiblast data at HH5 showing predicted trajectories. (C) Feature plots of selected genes across HH5 whole epiblast data. (D) Feature plots of selected genes across HH7 whole epiblast data. (E) UMAP embedding of subclusters from HH5 ectoderm cluster (HH5-Cl4) showing predicted trajectories. (F) UMAP embedding of clusters from all combined data set showing stages. (G) Feature plots of placode markers across all stages combined data set. (H) Feature plots of other selected genes across all stages combined data set. (I) Phase portraits (top row), velocity plots (middle row) and expression plots (bottom row) of selected genes across the HH7/ premigratory NC combined data sets.

## References

Azambuja, A. P. & Simoes-Costa, M. 2021. A regulatory sub-circuit downstream of Wnt signaling controls developmental transitions in neural crest formation. PLoS Genet, 17, e1009296.

Basch, M. L., Bronner-Fraser, M. & Garcia-Castro, M. I. 2006. Specification of the neural crest occurs during gastrulation and requires Pax7. Nature, 441, 218–22.

Bergen, V., Lange, M., Peidli, S., Wolf, F. A. & Theis, F. J. 2020. generalising RNA velocity to transient cell states through dynamical modeling. Nat Biotechnol, 38, 1408–1414.

Butler Tjaden, N. E. & Trainor, P. A. 2013. The developmental etiology and pathogenesis of Hirschsprung disease. Transl Res, 162, 1–15.

Chapman, S. C., Brown, R., Lees, L., Schoenwolf, G. C. & Lumsden, A. 2004. Expression analysis of chick Wnt and frizzled genes and selected inhibitors in early chick patterning. Dev Dyn, 229, 668–76.

Choi, H. M. T., Schwarzkopf, M., Fornace, M. E., Acharya, A., Artavanis, G., Stegmaier, J., Cunha, A. & Pierce, N. A. 2018. Third-generation in situ hybridization chain reaction: multiplexed, quantitative, sensitive, versatile, robust. Development, 145.

De Croze, N., Maczkowiak, F. & Monsoro-Burq, A. H. 2011. Reiterative AP2a activity controls sequential steps in the neural crest gene regulatory network. Proc Natl Acad Sci U S A, 108, 155–60.

Ezin, A. M., Fraser, S. E. & Bronner-Fraser, M. 2009. Fate map and morphogenesis of presumptive neural crest and dorsal neural tube. Dev Biol, 330, 221–36.

Fakhouri, W. D., Metwalli, K., Naji, A., Bakhiet, S., Quispe-Salcedo, A., Nitschke, L., Kousa, Y. A. & Schutte, B. C. 2017. Intercellular Genetic Interaction Between Irf6 and Twist1 during Craniofacial Development. Sci Rep, 7, 7129.

Gandhi, S., Ezin, M. & Bronner, M. E. 2020. Reprogrammeming Axial Level Identity to Rescue Neural-Crest-Related Congenital Heart Defects. Dev Cell, 53, 300–315 e4.

Hamburger, V. & Hamilton, H. L. 1951. A series of normal stages in the development of the chick embryo. J Morphol, 88, 49–92.

Hang, Y. & Stein, R. 2011. MafA and MafB activity in pancreatic beta cells. Trends Endocrinol Metab, 22, 364–73.

Haro, E., Delgado, I., Junco, M., Yamada, Y., Mansouri, A., Oberg, K. C. & Ros, M. A. 2014. Sp6 and Sp8 transcription factors control AER formation and dorsal-ventral patterning in limb development. PLoS Genet, 10, e1004468.

Hong, C. S. & Saint-Jeannet, J. P. 2017. Znf703, a novel target of Pax3 and Zic1, regulates hindbrain and neural crest development in Xenopus. Genesis, 55.

Ingraham, C. R., Kinoshita, A., Kondo, S., Yang, B., Sajan, S., Trout, K. J., Malik, M. I., Dun-Nwald, M., Goudy, S. L., Lovett, M., Murray, J. C. & Schutte, B. C. 2006. Abnormal skin, limb and craniofacial morphogenesis in mice deficient for interferon regulatory factor 6 (Irf6). Nat Genet, 38, 1335–40.

Janesick, A., Tang, W., Ampig, K. & Blumberg, B. 2019. Znf703 is a novel RA target in the neural plate border. Sci Rep, 9, 8275.

Kasberg, A. D., Brunskill, E. W. & Steven Potter, S. 2013. SP8 regulates signaling centers during craniofacial development. Dev Biol, 381, 312–23.

Kawakami, Y., Esteban, C. R., Matsui, T., Rodriguez-Leon, J., Kato, S. & Izpisua Belmonte, J. C. 2004. Sp8 and Sp9, two closely related buttonhead-like transcription factors, regulate Fgf8 expression and limb outgrowth in vertebrate embryos. Development, 131, 4763–74.

Khudyakov, J. & Bronner-Fraser, M. 2009. Comprehensive spatiotemporal analysis of early chick neural crest network genes. Dev Dyn, 238, 716–23.

Kousa, Y. A., Zhu, H., Fakhouri, W. D., Lei, Y., Kinoshita, A., Roushangar, R. R., Patel, N. K., Agopian, A. J., Yang, W., Leslie, E. J., Busch, T. D., Mansour, T. A., Li, X., Smith, A. L., Li, E. B., Sharma, D. B., Williams, T. J., Chai, Y., Amendt, B. A., Liao, E. C., Mitchell, L. E., Bassuk, A. G., Gregory, S., Ashley-Koch, A., Shaw, G. M., Finnell, R. H. & Schutte, B. C. 2019. The TFAP2A-IRF6-GRHL3 genetic pathway is conserved in neurulation. Hum Mol Genet, 28, 1726–1737.

La Manno, G., Soldatov, R., Zeisel, A., Braun, E., Hochgerner, H., Petukhov, V., Lidschreiber, K., Kastriti, M. E., Lonnerberg, P., Furlan, A., Fan, J., Borm, L. E., Liu, Z., Van Bruggen, D., Guo, J., He, X., Barker, R., Sundstrom, E., Castelo-Branco, G., Cramer, P., Adameyko, I., Linnars-Son, S. & Kharchenko, P. V. 2018. RNA velocity of single cells. Nature, 560, 494–498.

Lecoin, L., Sii-Felice, K., Pouponnot, C., Eychene, A. & Felder-Schmittbuhl, M. P. 2004. Comparison of maf gene expression patterns during chick embryo development. Gene Expr Patterns, 4, 35–46.

Lee, B. K., Shen, W., Lee, J., Rhee, C., Chung, H., Kim, K. Y., Park, I. H. & Kim, J. 2015. Tgif1 Counterbalances the Activity of Core Pluripotency Factors in Mouse Embryonic Stem Cells. Cell Rep, 13, 52–60.

Leung, A. W., Murdoch, B., Salem, A. F., Prasad, M. S., Gomez, G. A. & Garcia-Castro, M. I. 2016. WNT/beta-catenin signaling mediates human neural crest induction via a pre-neural border intermediate. Development, 143, 398–410.

Li, B., Kuriyama, S., Moreno, M. & Mayor, R. 2009. The posteriorizing gene Gbx2 is a direct target of Wnt signalling and the earliest factor in neural crest induction. Development, 136, 3267–78.

Liu, W., Lagutin, O. V., Mende, M., Streit, A. & Oliver, G. 2006. Six3 activation of Pax6 expression is essential for mammalian lens induction and specification. EMBO J, 25, 5383–95.

Lunn, J. S., Fishwick, K. J., Halley, P. A. & Storey, K. G. 2007. A spatial and temporal map of FGF/Erk1/2 activity and response repertoires in the early chick embryo. Dev Biol, 302, 536–52.

Pauli, S., Bajpai, R. & Borchers, A. 2017. CHARGEd with neural crest defects. Am J Med Genet C Semin Med Genet, 175, 478–486.

Prasad, M. S., Uribe-Querol, E., Marquez, J., Vadasz, S., Yardley, N., Shelar, P. B., Charney, R. M. & Garcia-Castro, M. I. 2020. Blastula stage specification of avian neural crest. Dev Biol, 458, 64–74.

Roellig, D., Tan-Cabugao, J., Esaian, S. & Bronner, M. E. 2017. Dynamic transcriptional signature and cell fate analysis reveals plasticity of individual neural plate border cells. Elife, 6.

Sahara, S., Kawakami, Y., Izpisua Belmonte, J. C. & O’leary, D. D. 2007. Sp8 exhibits reciprocal induction with Fgf8 but has an opposing effect on anterior-posterior cortical area patterning. Neural Dev, 2, 10.

Schille, C. & Schambony, A. 2017. Signaling pathways and tissue interactions in neural plate border formation. Neurogenesis (Austin), 4, e1292783.

Schlosser, G. 2008. Do vertebrate neural crest and cranial placodes have a common evolutionary origin? Bioes-says, 30, 659–72.

Siismets, E. M. & Hatch, N. E. 2020. Cranial Neural Crest Cells and Their Role in the Pathogenesis of Craniofacial Anomalies and Coronal Craniosynostosis. J Dev Biol, 8.

Streit, A. 2002. Extensive cell movements accompany formation of the otic placode. Dev Biol, 249, 237–54.

Stuart, T., Butler, A., Hoffman, P., Hafemeister, C., Papalexi, E., Mauck, W. M., 3RD, Hao, Y., Stoeckius, M., Smibert, P. & Satija, R. 2019. Comprehensive Integration of Single-Cell Data. Cell, 177, 1888–1902 e21.

Tomolonis, J. A., Agarwal, S. & Shohet, J. M. 2018. Neuroblastoma pathogenesis: deregulation of embryonic neural crest development. Cell Tissue Res, 372, 245–262.

Vega-Lopez, G. A., Cerrizuela, S., Tribulo, C. & Aybar, M. J. 2018. Neurocristopathies: New insights 150 years after the neural crest discovery. Dev Biol, 444 Suppl 1, S110–S143.

Wang, X., Liu, J., Zhang, H., Xiao, M., Li, J., Yang, C., Lin, X., Wu, Z., Hu, L. & Kong, X. 2003. Novel mutations in the IRF6 gene for Van der Woude syndrome. Hum Genet, 113, 382–6.

Williams, R. M., Candido-Ferreira, I., Repapi, E., Gavriouchkina, D., Senanayake, U., Ling, I. T. C., Telenius, J., Taylor, S., Hughes, J. & Sauka-Spengler, T. 2019. Reconstruction of the Global Neural Crest Gene Regulatory Network In Vivo. Dev Cell, 51, 255–276 e7.

Williams, R. M. & Sauka-Spengler, T. 2021a. Dissociation of chick embryonic tissue for FACS and preparation of isolated cells for genome-wide downstream assays. STAR Protoc, 2, 100414.

Williams, R. M. & Sauka-Spengler, T. 2021b. ex ovo electroporation of early chicken embryos. STAR Protoc, 2, 100424.

Yang, X., Chrisman, H. & Weijer, C. J. 2008. PDGF signalling controls the migration of mesoderm cells during chick gastrulation by regulating N-cadherin expression. Development, 135, 3521–30.

Ye, J. & Blelloch, R. 2014. Regulation of pluripotency by RNA binding proteins. Cell Stem Cell, 15, 271–280.

Zembrzycki, A., Griesel, G., Stoykova, A. & Mansouri, A. 2007. Genetic interplay between the transcription factors Sp8 and Emx2 in the patterning of the forebrain. Neural Dev, 2, 8.

Zhang, J., Tam, W. L., Tong, G. Q., Wu, Q., Chan, H. Y., Soh, B. S., Lou, Y., Yang, J., Ma, Y., Chai, L., Ng, H. H., Lufkin, T., Robson, P. & Lim, B. 2006. Sall4 modulates embryonic stem cell pluripotency and early embryonic development by the transcriptional regulation of Pou5f1. Nat Cell Biol, 8, 1114–23.

Zhang, S., Li, J., Lea, R., Vleminckx, K. & Amaya, E. 2014. Fezf2 promotes neuronal differentiation through localised activation of Wnt/beta-catenin signalling during forebrain development. Development, 141, 4794–805.

Zheng, G. X., Terry, J. M., Belgrader, P., Ryvkin, P., Bent, Z. W., Wilson, R., Ziraldo, S. B., Wheeler, T. D., Mcdermott, G. P., Zhu, J., Gregory, M. T., Shuga, J., Montesclaros, L., Under-Wood, J. G., Masquelier, D. A., Nishimura, S. Y., Schnall-Levin, M., Wyatt, P. W., Hindson, C. M., Bharadwaj, R., Wong, A., Ness, K. D., Beppu, L. W., Deeg, H. J., Mcfarland, C., Loeb, K. R., Valente, W. J., Ericson, N. G., Stevens, E. A., Radich, J. P., Mikkelsen, T. S., Hindson, B. J. & Bielas, J. H. 2017. Massively parallel digital transcriptional profiling of single cells. Nat Commun, 8, 14049.

